# Pushing the limits: expanding the temperature tolerance of a coral photosymbiont through differing selection regimes

**DOI:** 10.1101/2024.02.11.579409

**Authors:** Hugo J. Scharfenstein, Lesa M. Peplow, Carlos Alvarez-Roa, Matthew R. Nitschke, Wing Yan Chan, Patrick Buerger, Madeleine JH van Oppen

## Abstract

Coral thermal bleaching resilience can be improved by enhancing photosymbiont thermal tolerance via experimental evolution. While successful for some strains, selection under stable temperatures was ineffective at increasing the thermal threshold of an already thermo-tolerant photosymbiont (*Durusdinium trenchii*). Corals from environments with fluctuating temperatures tend to have comparatively high heat tolerance. Therefore, we investigated whether exposure to temperature oscillations can raise the upper thermal limit of *D. trenchii*. We exposed a *D. trenchii* strain to stable and fluctuating temperatures profiles, which varied in oscillation frequency. After 2.1 years (54-73 generations), we characterised the adaptive responses under the various experimental evolution treatments by constructing thermal performance curves of growth from 21 to 31°C for the heat-evolved and wild-types lineages. Additionally, oxidative stress, photophysiology, photosynthesis and respiration rates were assessed under increasing temperatures. Of the fluctuating temperature profiles investigated, selection under the most frequent oscillations (diurnal) induced the greatest widening of *D. trenchii*’s thermal niche. Continuous selection under elevated temperatures induced the only increase in thermal optimum and a degree of generalism. Our findings demonstrate how differing levels of thermal homogeneity during selection drive unique adaptative responses to heat in a coral photosymbiont.

## Introduction

Reef-forming corals (Scleractinia) are experiencing longer and more frequent marine heatwaves (Oliver *et al*., 2018), which are projected to gain in frequency, intensity and duration as a consequence of climate change (Oliver *et al*., 2019). Primarily driven by worsening marine heatwaves (Hughes *et al*., 2018), global coral cover has halved since the 1950s (Eddy *et al*., 2021). Under business-as-usual scenarios, the combined effects of climate change and local anthropogenic pressures could lead to 75% of the remaining coral reefs facing unsuitable environmental conditions as soon as 2035 (Setter *et al*., 2022).

To avert the decline of coral reefs, new management tools are urgently needed to enhance the capacity of corals to adapt to shifting climactic baselines (Bay *et al*., 2023). One proposed strategy is the development of coral stocks possessing enhanced thermal tolerance through (human)-assisted evolution (AE) (van Oppen *et al*., 2015). AE interventions include a range of host-centric (selective breeding, acclimatisation, translocation) and microbial manipulation methods, which are actively being explored by a growing number of coral reef restoration research and development initiatives (e.g., <u>RRAP</u>, <u>xReefs</u>, <u>CORDAP</u>). Promisingly, some studies applying AE strategies/principles have been able to demonstrate increases in coral thermal tolerance (Buerger *et al*., 2020; Chan *et al*., 2018, 2023; Drury *et al*., 2022; Quigley *et al*., 2020, 2023; Rosado *et al*., 2018).

The manipulation of the photosymbiont (Symbiodiniaceae) communities of corals is an AE intervention founded on knowledge that the thermal tolerance of Symbiodiniaceae strongly influences the susceptibility of corals to withstand thermal stress (reviewed by van Oppen & Nitschke, 2022). Natural variation in thermal tolerance occurs among Symbiodiniaceae, with differences of 1-2°C in thermal tolerance reported between *Cladocopium* spp. and *Durusdinium* spp. (i.e., two predominant photosymbiont genera in Indo-Pacific corals) (Abrego *et al*., 2008; Berkelmans & Van Oppen, 2006; Silverstein *et al*., 2017). However, algal symbiont manipulation with naturally-occurring thermally tolerant lineages is constrained by two factors: 1) growth trade-offs have been documented in corals hosting the thermally-tolerant *Durusdinium* strains (Cantin *et al*., 2009; Little *et al*., 2004), though not in all parts of the world or taxa (Turnham *et al*., 2023); and 2) other thermally-tolerant Symbiodiniaceae (e.g., *Fugacium* spp.) rarely associate with corals (LaJeunesse *et al*., 2003), though see (Terraneo *et al*., 2023).

To circumvent some of these limitations, experimental evolution of Symbiodiniaceae via thermal selection can generate strains that confer high temperature tolerance, while maintaining coral growth rates (Chan *et al*., 2023). Experimental evolution involves the propagation of an organism under controlled conditions, and this approach has historically been applied to microorganisms to study evolutionary processes (reviewed by Kawecki *et al*., 2012). In the context of coral reef restoration, experimental evolution has been used to generate Symbiodiniaceae lineages with increased thermal tolerance (Chakravarti *et al*., 2017; Chakravarti & van Oppen, 2018; Scharfenstein *et al*., 2023). Following symbiosis establishment with corals, some heat-evolved *Cladocopium proliferum* lineages were found to increase the thermal bleaching resilience of their host to levels comparable to naturally-occurring *Durusdinium* symbionts, all whilst not incurring the expected trade-off in growth (Chan *et al*., 2023).

To date, thermal selection of Symbiodiniaceae has relied exclusively on exposure to stable, elevated temperatures, typically ranging from 30-33°C (Chakravarti *et al*., 2017; Chakravarti & van Oppen, 2018; Huertas *et al*., 2011; Pierangelini *et al*., 2020; Scharfenstein *et al*., 2023). Occasionally, temperatures were ratcheted upwards throughout the selection process after observing positive growth over a three to six week period at a specific temperature. With the exception of two *Durusdinium* strains (SCF082.01 and SCF086.01) (Chakravarti & van Oppen, 2018; Scharfenstein *et al*., 2023), all heat-evolved symbionts responded positively to continuous thermal selection, displaying a stable adaptive change to thermal stress. Increased thermal tolerance was typically characterised by faster *in vitro* growth rates and, in some cases, higher photosynthetic efficiencies in the selected Symbiodiniaceae over their wild-type counterparts (Buerger *et al*., 2020, 2023; Chakravarti *et al*., 2017; Chakravarti & van Oppen, 2018; Scharfenstein *et al*., 2023).

Over the last decade, evidence has accumulated that corals inhabiting reef environments characterized by strong environmental variability (e.g., diel temperature/pH fluctuations in tidal pools and mangrove systems) display enhanced stress tolerance over corals living under comparatively more stable conditions (Camp *et al*., 2017; Oliver & Palumbi, 2011; Palumbi *et al*., 2014; Schoepf *et al*., 2015, 2020). Recent experimental evolution studies on marine microalgae have also shown that fluctuating conditions can increase thermal tolerance (reviewed by Chan *et al*., 2021). Notably, selection of a marine diatom under fluctuating temperatures has been linked to faster adaptation compared to selection under continuous exposure to elevated temperatures (Schaum *et al*., 2018).

Coastal sea surface temperatures tend to fluctuate substantially on a diurnal basis and more broadly on a seasonal timescale, punctuated by the sporadic occurrences of marine cold-spells and heatwaves. Highly frequent temperature fluctuations (i.e., on a diurnal basis) have been shown to promote coral thermal tolerance (Safaie *et al*., 2018; Schoepf *et al*., 2022), though other studies reported no (Schoepf *et al*., 2019) or detrimental (Klepac & Barshis, 2020) effects. Less frequent temperature fluctuations (i.e., spanning days to weeks) have also been shown to induce protection from subsequent thermal stress events, though the pattern of the frequency can lead to contrasting outcomes (Ainsworth *et al*., 2016). Beyond species-specific mechanisms that may influence adaptation under thermal fluctuations, we lack a mechanistic understanding of how temperature fluctuations influence adaptation (Schoepf *et al*., 2022). Adaptation is likely influenced by the stochasticity (Gill *et al*., 2022), the magnitude and the frequency (Schoepf *et al*., 2022) of the fluctuation in relation to the organism’s generation time. Intra- and inter-generational fluctuations (e.g., diurnal vs. weekly fluctuations) could favour differing algal symbiont genotypes (Chan *et al*., 2021). To the best of our knowledge, no study has investigated the evolutionary outcomes of these differing frequencies of temperature fluctuations for Symbiodiniaceae.

In this study, we aimed to determine whether periodic, rather than continuous, thermal selection may yield greater increases in thermal tolerance of Symbiodiniaceae (i.e., increased temperature limits of growth, *sensu* Barton *et al*., 2023) and, if so, how the frequency of fluctuations may influence adaptation. A *Durusdinium trenchii* strain (SCF086.01) that previously responded poorly to continuous experimental evolution under elevated temperature (Chakravarti & van Oppen, 2018) was subjected to three different fluctuating temperature profiles: fluctuations between ambient and elevated temperatures on a diurnal basis, every three weeks (corresponding to 2-3 generations) and every six weeks (4-5 generations). Cultures were also continuously exposed to either elevated or ambient temperatures. Following 2.1 years of experimental evolution (54-73 generations), traits relating to growth, photosynthesis and oxidative stress of the heat-evolved and ancestral lineages were characterized across their natural range of temperatures.

## Materials and methods

### Organism and culture conditions

The heterogenous (polyclonal) *Durusdinium trenchii* culture (SCF086.01, ITS2 type profile: D1-D4-D4c-D1l-D6-D4f-D4au-D1c-D1h-D1ih-D6c-D1hf) used in this study was isolated and provided by the Symbiont Culture Facility at the Australian Institute of Marine Science, Australia. The strain was isolated in November 2011 from a *Porites lobata* colony which originated from Davies Reef, Australia, and has since been maintained at 27°C.

During the subsequent experiments, sub-culturing was performed by transferring biomass to 0.2 μm filter-sterilized seawater with added Daigo’s IMK Medium (Nihon Pharmaceutical Co. Ltd., Tokyo, JP, hereafter ‘IMK’). The *D. trenchii* cultures were maintained in environmental chambers (Steridium, Brendale, AU), which followed a 14 h:10 h light:dark cycle and where the ambient light intensity was set to 60 ± 10 µmol/m^2^/s (Sylvania F15W/T8/865 light tubes; Sylvania, Newhaven, UK).

### Experimental evolution

In November 2020, *D. trenchii* biomass growing at 27°C was sub-cultured into 30 cell culture flasks (25cm^2^, 0.2 μm vent cap; Sigma-Aldrich, St. Louis, MO, US). Throughout experimental evolution sub-culturing was performed by transferring biomass into a new flask to achieve 200,000 cells/ml in 20 mL of IMK. The cultures were then equally distributed to five temperature profiles (TPs) (see Fig. 1 for illustration of TPs). From this point onwards, the cultures were maintained as independent lineages, with six replicates subjected to each TP. The TPs consisted of: diurnal fluctuations between ambient temperatures (during the 10 h dark period) and elevated temperatures (during the 14 h light period), hereafter ‘Fluc-short’; symmetrical fluctuations between ambient and elevated temperatures every third week, corresponding to 2-3 generations at each temperature, hereafter ‘Fluc-med’; asymmetrical fluctuations between temperatures with ambient maintained for three weeks and elevated for six weeks, which corresponds to 4-5 generations, hereafter ‘Fluc-long’; continuous exposure to elevated, hereafter ‘Cont-ele’; and finally, continuous exposure to ambient temperatures, hereafter ‘Cont-amb’ (see Fig. S1 for additional detail on the TPs).

**Figure 1:**
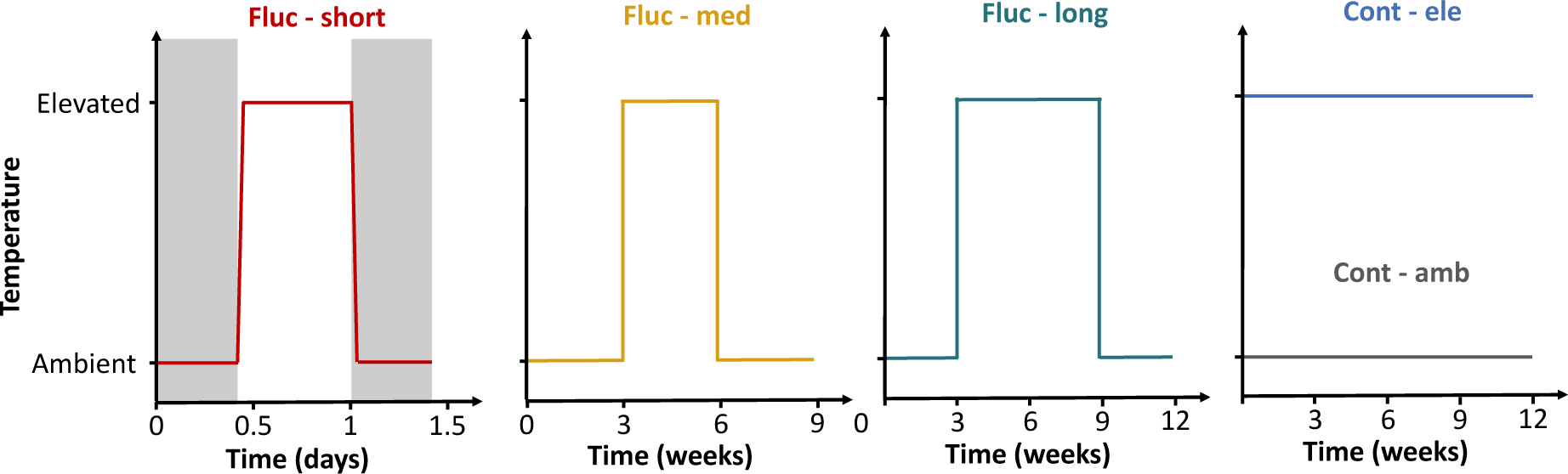
Diagram of temperature profiles used during experimental evolution. For Fluc-short, shaded areas correspond to the dark phase, whilst white areas correspond to the light phase. Ramping between temperature levels was performed over one hour. With the exception of Cont-amb, the elevated temperature of the temperature profiles was ratcheted upwards by 1°C according to the criterium defined in Materials and Methods.

Except for cultures subjected to Cont-amb (which acted as a control profile), the elevated temperature of the TPs was ratcheted up by 1°C. Starting from 28°C, ratcheting was carried out when the following criterion was met: mean growth rates from all six lineages exposed to the same TP needed to be greater or equal to those of cultures from Cont-amb over a six week period. After 43 weeks, the condition for ratcheting was relaxed due to it only enabling the ratcheting of elevated temperatures in Fluc-short. For the remainder of the experimental evolution, the criterion was changed to: mean growth rates from all six lineages exposed to the same TP needed to be positive over a six week period (according to Chakravarti *et al*., 2017).

To avoid cultures entering a nutrient depleted state that would limit growth rates, sub-culturing was carried out every third to fourth week, matching the temperature cycles of Fluc-med and Fluc-long. Prior to sub-culturing, cell densities of the cultures were measured via flow cytometry (C6 Plus Flow Cytometer, BD Biosciences, Franklin Lakes, NJ, US) according to Scharfenstein *et al*. (2023). Growth rates were calculated for each lineage using the following equation:

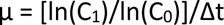

where C_1_/C_0_ are the cell densities at the end/start of the growth period and Δt is the duration of the growth period in days.

If negative growth rates were recorded for all six lineages from the same TP (i.e., all cultures were crashing in response to the elevated temperature), then the cultures were moved back to 27°C to recover until positive growth was measured across all six lineages. These were then subjected once again to the TP.

### Thermal performance assay

#### Experimental design

After 110 weeks (2.1 years) of experimental evolution (see Fig. S2 for lineage specific number of generations), physiological traits of all 30 *D. trenchii* lineages were assessed across a range of temperatures. The aim of this ‘thermal performance assay’ was to quantify adaptive responses from long-term exposure to the differing thermal selection strategies.

Biomass from each lineage was transferred to 24-well plates (TPP tissue culture plates; TPP, Trasadingen, CH). The cultures were grown at 27°C for six weeks to pre-acclimate all cultures to the same conditions. During this period, sub-culturing was carried out every two weeks to grow sufficient biomass. The cultures were then ramped down/up at a rate of 2°C/day to temperatures spanning 21-31°C. This temperature range was chosen as it broadly encompasses the annual temperatures the *D. trenchii* strain would have encountered at its site of origin (see Fig. S3 for the temperature profile of Davies Reef).

Once the target temperature was reached, the lineages were sub-cultured into fresh IMK culture medium at a cell density of 100,000 cells/ml and culture volume of 2 ml. Lineages originating from different TPs were kept in different 24-well plates to avoid any cross-contamination between experimental evolution treatments during the thermal performance assay. At each temperature, biomass from each lineage was distributed across four wells in each plate (n_tech_ = 4).

Culture growth was monitored across temperatures every third-fourth day, except for cultures at 31°C, which were measured every second day, with measurements alternating between two replicates. The more frequent sampling regime at 31°C was to ensure sufficient timepoints would be obtained to model growth curves before the cultures crashing. Once the cultures were deemed to be in their exponential growth phases (see Fig. S4 for growth curves), biomass from each lineage was sub-cultured to a new plate in fresh IMK following the aforementioned procedure. Unless specified otherwise, the reported measurements (i.e., photochemical efficiency, oxidative stress, photosynthesis and respiration rates) from the thermal performance assay were taken during the exponential phase of this second growth cycle (see Fig. S4) in order to minimize the influence of acclimation in our assessment of stable adaptation to the growth temperatures. Growth rates were also derived from the second growth cycle.

#### Growth rate measurements

One hour after the onset of the photoperiod, the optical density (OD) of each lineage at each temperature was measured using a microplate reader (Synergy H4, Agilent BioTek, Santa Clara, US) at 670 nm. Immediately before OD measurements, all cultures were manually resuspended in each well through pipetting. Measurements were repeated on the same wells every three-four days, on average. For each lineage at each temperature, the median OD of all four well (technical) replicates was calculated. The OD values of empty wells containing IMK, which were kept at the same growth temperature and measured on the same day the lineages, were then subtracted from the culture OD.

OD measurements were converted to cell densities using calibration curves, which were established at each growth temperature prior to the start of the experiment (see Fig. S5 for calibration curves). Growth rates were calculated by fitting parametric non-linear models to the growth curves using the R package *nls.multstart* (version 1.2.0) and following a modified procedure from Padfield *et al*. (2019). In brief, six different growth models, derived from Gompertz, Baranyi and Buchanan models (obtained from the R package *nlsMicrobio*; version 0.0-3) were fitted to the growth data. Three of the models (*baranyi, baranyi_no_*lag, *gompertz*) were rejected as they provided erroneous maximum growth rate (µ_max_) estimates, particularly at the extreme temperatures (see Fig. S6 for parameter estimates). The remaining models (*buchanan*, *buchanan_no_lag*, *baranyi_no_K*) were then filtered for the best fit. For each lineage at each temperature, models with a small sample-size corrected Akaike Information Criterion (AICc) score within Δ2 AICc of the fit with lowest AICc score were kept. Out of the remaining models, the model that provided the lowest confidence interval for the parameter µ_max_ was kept. Following this filtering process, the *buchanan_no_lag* model was found to be the best fit in 149/180 cases and selected as final model to derive µ_max_.

#### Thermal performance curve fitting

Thermal performance curves (TPCs) of previously calculated maximum growth rates (µ_max_) were constructed in R using the package *rTPC* (version 1.0.2; Padfield *et al*., 2021). Five nonlinear regression models (*thomas_2012, thomas_2017*, *joehnk_2008, lactin2_1995, kamykowski_1985*) were selected based on their ability to account for negative growth rates and have been successfully applied in constructing TPCs with marine phytoplankton (Barton *et al*., 2023). Other models (e.g., Gaussian, Beta) previously used to construct TPCs for Symbiodiniaceae (Dilernia *et al*., 2023) and corals (Álvarez-Noriega *et al*., 2023; Jurriaans & Hoogenboom, 2019) were considered but not used since they did not allow for the fitting of negative growth rates. Models were fitted to each lineage and incorporated a model weight corresponding to the standard deviation in µ_max_ observed for each experimental evolution treatment at each growth temperature.

The models *thomas_2017*, *kamykowski_1985* and *joehnk_2008* were found to provide the lowest AICc score in a roughly equal number of cases (11/30, 11/30 and 8/30 cases, respectively). No single model was found to provide the best fit across one experimental evolution treatment. To avoid introducing a bias from the selection of one model over another, we selected the three aforementioned models and proceeded with the averaging of models that were within Δ2 AICc of the best model (see Fig. S7-S8 for TPC fits). The following parameters of interest were subsequently extracted from TPCs fitted with the model average: the thermal optimum (T_opt_; i.e., temperature at which the highest µ_max_ is recorded), the thermal breadth (T_br_; temperature range at which the µ_max_ is at least 80% of r_max_), the thermal safety margin (TSM; temperature difference between CT_max_ and T_opt_), the maximum rate (r_max_; the highest µ_max_ of the TPC), the critical thermal maximum (CT_max_; upper temperature where the predicted µ_max_ is closest to 0) and minimum (CT_min_).

#### Photosynthesis and respiration measurements

Oxygen (O_2_) consumption and production was measured using an optical O_2_ meter (FireSting-Pro; PyroScience, Aachen, DE) connected to O_2_ sensor spots (OXSP5-ADH; PyroScience) mounted on custom-built air-tight chambers. Due to only 12 spots and channels being available, measurements were carried out in three consecutive runs between 5 and 8 hours into the photoperiod (i.e., one run = 12 samples). For each run, we ensured that two lineages from each TP were present. The lineages were measured in the same order between growth temperatures to control for time of day. Prior to measurements, cultures were centrifuged (3,000 g x 5 min) to partly separate the Symbiodiniaceae from smaller prokaryotic cells (according to Pierangelini *et al*., 2020). The supernatant was kept as a blank for subsequent measurements to account for remaining loosely associated bacteria. The pellet was resuspended in fresh IMK (pre-exposed for 1-2 hours to the growth temperature) at a normalized density of 750,000 cells/ml. The samples were then returned to their growth temperature for two hours for temperatures of the culture medium to stabilize. Prior to transferring 1.46 ml of the normalized biomass to the chambers, cultures were vortexed and shaken to ensure O_2_ saturation in the culture medium. Once the samples were transferred, the chambers were gas-tight sealed with an O-ring while ensuring no air bubbles remained.

The cultures were then placed in a temperature-controlled incubator and dark-adapted for 15 min, after which the O_2_ consumption was measured for 5 mins to determine dark respiration (R_d_) rates. Following the dark period, lights were turned on (at growth irradiance, i.e., 60 ± 10 µmol/m^2^/s) and O_2_ production was recorded over 25 min. Net photosynthesis (P_net_) rates were measured over 5 mins in a sliding window corresponding to 5-15 mins after the start of the light period. Blanks were included to account for any prokaryotic contribution to R_d_ and P_net_. Raw O_2_ measurements are available in Fig. S9. During measurements, we ensured the cultures were homogenized using an orbital shaker (2 rotations per second). Measurements were carried out at the same time of day between temperatures.

R_d_ and P_net_ rates were calculated using the R package *respR* (version 2.2.0; Harianto *et al*., 2019). Raw O_2_ measurements were first visually inspected for irregularities (i.e., highly variables rates within the timeframes used to determine R_d_ and P_net_). For some samples, O_2_ production during the light phase window was found to be non-linear (corresponding to: 1 lineage per treatment at 31°C, 2 lineages for Cont-ele, Cont-amb and 1 for Fluc-short, Fluc-med at 29°C, 1 lineage for Fluc-med at 27°C). As a result, these samples were omitted from the calculation of P_net_ rates (see Fig. S9). Gross photosynthesis (P_gross_) was calculated as the sum of P_net_ and R_d_. For each sample, R_d_ and P_net_ rates were adjusted with the rates measured from the blank obtained for its respective experimental evolution treatment.

#### Oxidative stress

Oxidative stress was assessed by measuring the accumulation of extracellular reactive oxygen species (ExROS) in the culture medium of lineages grown at 27, 29 and 31°C according to Buerger *et al*. (2020) and Scharfenstein *et al*. (2023). Briefly, cultures were centrifuged at 15,000 g for 5 min, the supernatant removed and stained with CellROX Orange Reagent (Thermo Fisher Scientific, Waltham, MA, US) prior to fluorescence being measured with a microplate reader (Synergy H4). Three technical replicate measurements were carried out for each lineage, which were averaged prior to normalisation by cell density.

#### Photochemical efficiency

The photochemical efficiency of each lineage was measured using an imaging pulse-amplitude modulation chlorophyll fluorometer (Maxi version IMAGING PAM M-Series; Walz, Effeltrich, DE). Two hours after the end of dark period, cultures were dark-adapted at their respective temperature for 15 mins before being placed under the IMAGING PAM. The maximum quantum yield of photosystem II (PSII) of the dark-adapted cultures (F_v_/F_m_ = [F_m_-F_0_]/F_m_) was then measured. One area of interest encompassing the entirety of the well was set for each well. The F_v_/F_m_ for each lineage was calculate as the median F_v_/F_m_ of the four plate (technical) replicates. The following IMAGING PAM parameters were used: light intensity = 5, gain = 1, damping = 2, saturating pulse width = 0.8s, saturating pulse intensity = 10s.

#### Melting temperature of the photosynthetic membrane

The melting temperature (T_m_) of the thylakoid membrane of each lineage grown at 27, 29 and 31°C was measured following the methodology described by Díaz-Almeyda *et al*. (2011) and Beltrán *et al*. (2021). Biomass from each lineage was normalised to 750,000 cells/ml. The normalised samples were then aliquoted (200 µl per lineage) to three replicate 96-well plates. One plate was dark-adapted for 15 min at 27°C, transferred to a thermocycler (SuperCycler; KyraTec, Mansfield, AU) and incubated at 27°C for 5 min. At the end of the incubation period the F_v_/F_m_ of all the samples was measured using an IMAGING-PAM mounted above the thermocycler (using the same settings as above, with the exception of light intensity = 3). The temperature was then immediately ramped up to 33°C and the sample incubated for a further 5 min, after which the F_v_/F_m_ was measured. The incubation duration and F_v_/F_m_ measurements were repeated iteratively by 1°C increments until 43°C. The F_v_/F_m_ values obtained from each temperature were then converted to values relative to the F_v_/F_m_ measurements at 27°C.

Using the R package *drc* (version 3.0-1; Ritz *et al*., 2015), data points were fitted with Weibull (*W1.4*), Logistic (*L.4*) and Gaussian (*G.4*) models. The model with the lowest AICc score was then kept, which overwhelmingly corresponded to the Weibull model (83/90 fits) (see Fig. S10 for fits). The effective dose at 50% (ED_50_) was calculated for each curve, which determined the T_m_ of the photosynthetic membrane for each lineage. Thermal plasticity of the thylakoid membrane was calculated as the difference between melting temperatures at 31 and 27°C, according to Mansour *et al*. (2018).

#### Statistical analyses

Statistical analyses were performed in R using the package *stats* (version 4.3.0.). To determine whether any significant shifts in traits resulted from the experimental evolution treatments, a one-way analysis of variance (ANOVA) followed by Tukey’s post-hoc test was used. Model assumptions of normal distribution and homogeneity of variance were verified using QQplots and boxplots. The following traits and parameters were tested for significant differences between experimental evolution treatments: TPC parameters (T_opt_, T_br_, TSM, r_max_, CT_min/max_), R_d_, P_net_, P_gross_, F_v_/F_m_, T_m_ and ExROS levels. Apart from the TPC parameters, were derived from growth cycles, analyses were performed with measurements collected from the exponential phase of the second growth cycle (i.e., no repeated measures over time).

## Results

### Experimental evolution

Exposure to 31°C had a strong negative effect on the growth of the heat-evolved *D. trenchii* lineages, regardless of the thermal selection regime (Fig. 2A; Fig. S11). Cultures from Fluc-short, Fluc-long and Cont-ele crashed following initial exposure to 31°C (i.e., negative growth rates were recorded). However, upon second exposure to 31°C lineages subjected to Fluc-short achieved positive growth, as opposed to cultures from Fluc-med and Fluc-long. Lineages from Fluc-short subsequently exhibited positive growth at 32°C (0.05 day^−1^, σ=0.008), displaying comparable growth rates to those observed at 31°C (0.05 day^−1^, σ=0.02). As a result, Fluc-short was eventually ratcheted up to 33°C, the highest elevated temperature of all selection regimes. Fluc-short was the only TP where cultures were viably maintained above 31°C.

**Figure 2:**
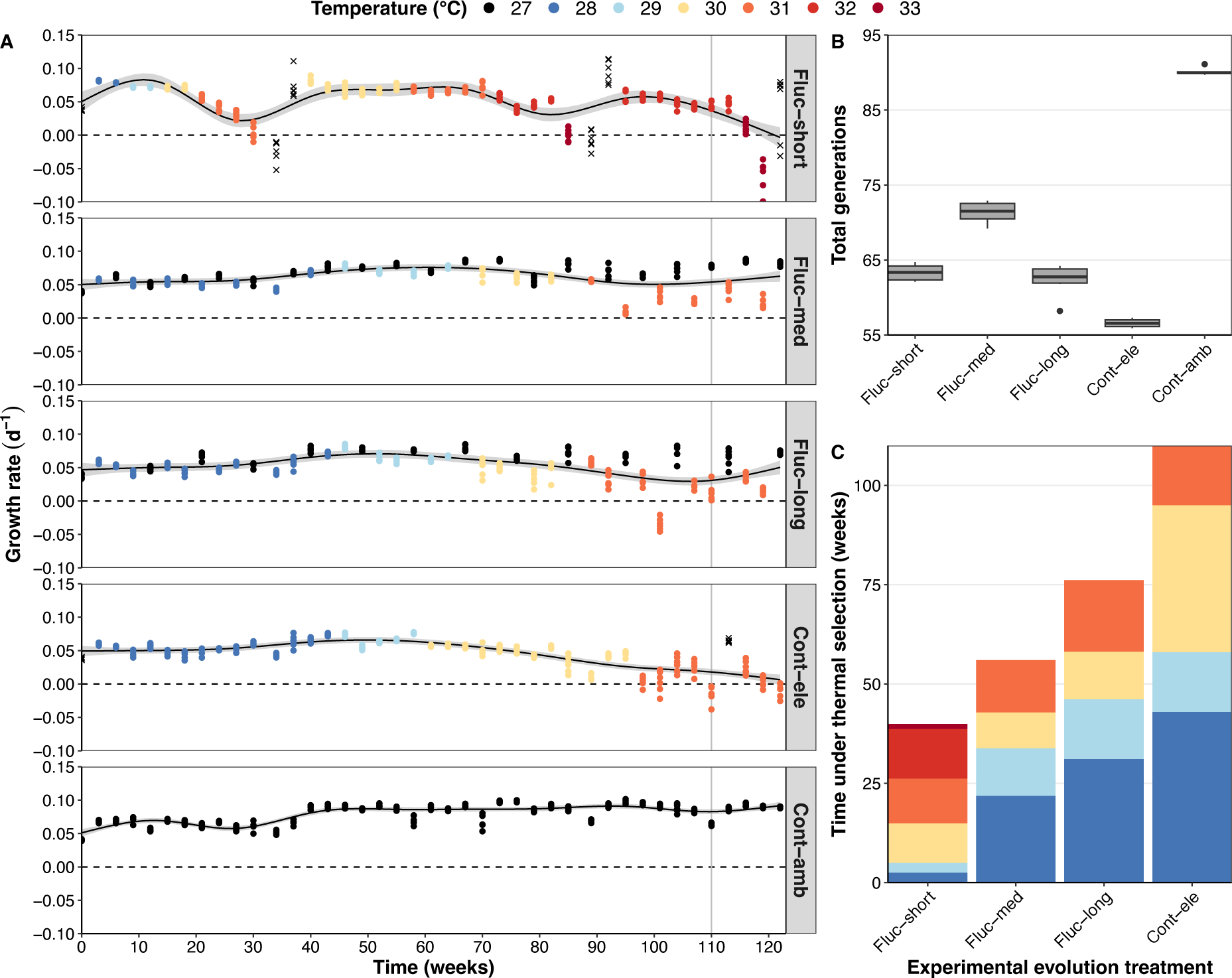
Growth rate trajectories of *Durusdinium trenchii* lineages (A), total number of generations elapsed (B) and time spent under thermal selection (C) during experimental evolution. In (**A**), colours indicate the temperature during the growth cycle (3-4 weeks). In the case of Fluc-short, colours represent the elevated temperature during that growth cycle, though lineages were exposed to both ambient (27°C) and elevated temperatures within one cycle. If negative growth rates were measured in response to the thermal selection regime (i.e., the cultures crashed), then the cultures were returned to ambient temperature (27°C) until positive growth rates were obtained, after which the cultures were subjected to elevated temperatures once again. Such unplanned relaxations of temperatures are designated by a cross (X). The vertical grey line indicates the start of the thermal performance assay. For (**A**) and (**B**), growth rates and generations were calculated off six lineages per temperature profile. For (**B**) and (**C**), number of generations and time under thermal selection were calculated till the start of the thermal performance assay. Experimental evolution treatments: Fluc-short = diurnal fluctuations, Fluc-med = fluctuations every 2-3 generations, Fluc-long = fluctuations every 4-5 generations, Cont-ele = continuous exposure to elevated temperatures, Cont-amb = continuous exposure to ambient temperature.

Cont-ele led to declining growth rates as temperatures increased (from 0.066 day^−1^ at 29°C to 0.008 day^−1^ at 31°C). As a result, only 54-57 generations elapsed during experimental evolution for Cont-ele lineages, the smallest evolutionary window out of any experimental evolution treatment (Fig. 2B). The periodic relaxation of temperatures meant higher growth rates could be maintained at elevated temperatures (e.g., 0.032 day^−1^ for Fluc-med and 0.022 day^−1^ for Fluc-long at 31°C). This translated to a higher number of generations (62-73) elapsing for Fluc-short, Fluc-med and Fluc-long lineages. However, comparatively to Cont-ele, fewer generations were spent at elevated temperature (Fig. S2).

Our initial criterion to ratchet an experimental evolution treatment to a higher temperature (i.e., higher growth rates in heat-selected than wild-type lineages from Cont-amb) led to Fluc-short being ratcheted to 31°C after only 21 weeks of experimental evolution (Fig. 2A). In contrast, Fluc-med, Fluc-long and Cont-ele were only ratcheted up once the criterion had been relaxed (i.e., positive growth over six weeks), only reaching 31°C after 89-98 weeks. Relative to the overall time spent at elevated temperature, these TPs were maintained longer below 31°C than Fluc-short, which was primarily maintained at 31-32°C (Fig. 2C).

### Thermal performance assay

#### Unique shifts in thermal performance according to selection strategy

Following 110 weeks of experimental evolution, the thermal performances of the heat-evolved and wild-type *D. trenchii* lineages were compared. TPCs revealed shifts in thermal performances that were unique to each thermal selection strategy (Fig. 3).

**Figure 3:**
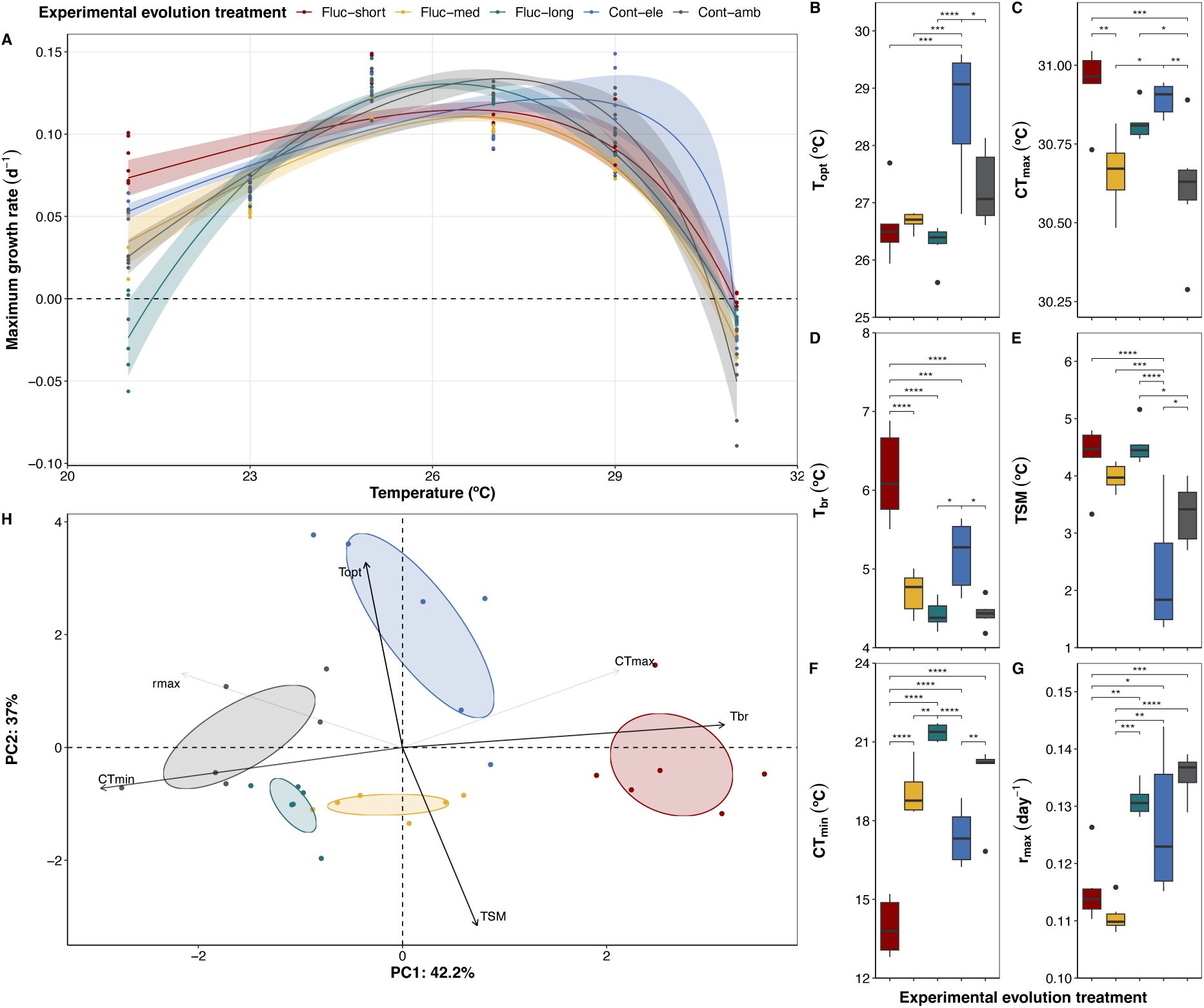
Thermal performance curves of maximum growth rates (A) and modelled parameters (B-G) of heat-evolved and wild-type *Durusdinium trenchii* lineages. A principal component analysis (H) was carried out with the TPC parameters for each experimental evolution treatment. (**A**) For visualization purposes, the mean TPC fit for each experimental evolution treatment is represented (lineage-specific TPC fits are available in Fig. S6). The ribbons represent the standard deviation across the TPCs for each treatment. (**B-G**) Modelled parameters were extracted from TPCs constructed for each lineage. T_opt_ = thermal optimum; CT_max_ = critical thermal maximum; T_br_ = thermal breadth; TSM = thermal safety margin; CT_min_ = critical thermal minimum; r_max_ = maximum growth rate. (**H**) The weight of each trait’s contribution to the PCA is represented by the shading of the arrows (darker = more contribution, light = less contribution). Ellipses reflect the 95% confidence intervals around the centroid of each cluster, which correspond to the experimental evolution treatments. Experimental evolution treatments: Fluc-short = diurnal fluctuations, Fluc-med = fluctuations every 2-3 generations, Fluc-long = fluctuations every 4-5 generations, Cont-ele = continuous exposure to elevated temperatures, Cont-amb = continuous exposure to ambient temperature. Levels of significance: * = p_adj_ < 0.05; ** = p_adj_ < 0.01; *** & **** = p_adj_ < 0.001.

Relative to Cont-amb, selection under Fluc-short significantly expanded the breadth of the *D. trenchii* thermal niche, characterised by a higher CT_max_ (+0.33°C) and lower CT_min_ (−6.1°C; Fig. 3A,C,F). Fluc-short lineages displayed the most significant increase in thermal tolerance (i.e., the upper temperature limit of growth, CT_max_). Cont-ele also induced an improvement of CT_max_ and CT_min_ over Cont-amb, leading to a significant increase in T_br_, albeit to a lesser extent than Fluc-short. In contrast, minimal shifts in TPC limits were observed for lineages selected under fluctuations spanning multiple generations (Fluc-med and Fluc-long), with no changes in T_br_ recorded.

Selection under fluctuating temperatures – regardless of frequency – did not change the T_opt_ of *D. trenchii* (Fig. 3B). Significant increases in the TSM of Fluc-short and Fluc-long lineages were recorded (+0.75-1.14°C relative to Cont-amb; Fig. 3E). In contrast, continuous thermal selection did lead to an increase in T_opt_, relative to all lineages, but decreased the TSM. The observed improvements in upper and lower thermal tolerance for Fluc-short lineages came with a trade-off against r_max_ (−0.02 day^-1^ relative to Cont-amb; Fig. 3G).

The TPCs revealed distinct thermal performances for all selective treatments (Fig. 3H). Changes in thermal performances of Fluc-short and Cont-ele lineages were driven primarily by shifts in T_br_ and T_opt_, respectively, suggesting differing adaptive trajectories in response to their respective thermal selection regimes.

#### Improved productivity associated with gains in thermal tolerance in Fluc-short

We next investigated photosynthesis and respiration rates of the heat-evolved lineages under increasing temperatures. None of the thermal selection strategies led to significant changes in the dark respiration (R_d_) rates of *D. trenchii,* regardless of temperature (Fig. 4A, Fig. S12). This suggests that experimental evolution did not alter the energetic burden of elevated temperature on *D. trenchii*. Increases in net photosynthesis (P_net_) rates were only found to be significant for Fluc-short and Fluc-long at 31°C (Fig. 4B, Fig. S12). When gross photosynthesis (P_gross_) is considered, Fluc-short lineages displayed the greatest improvement, relative to Cont-amb at 31°C (Fig. S12). No significant differences in P_net_ and P_gross_ were observed between Cont-ele and Cont-amb at 31°C.

**Figure 4:**
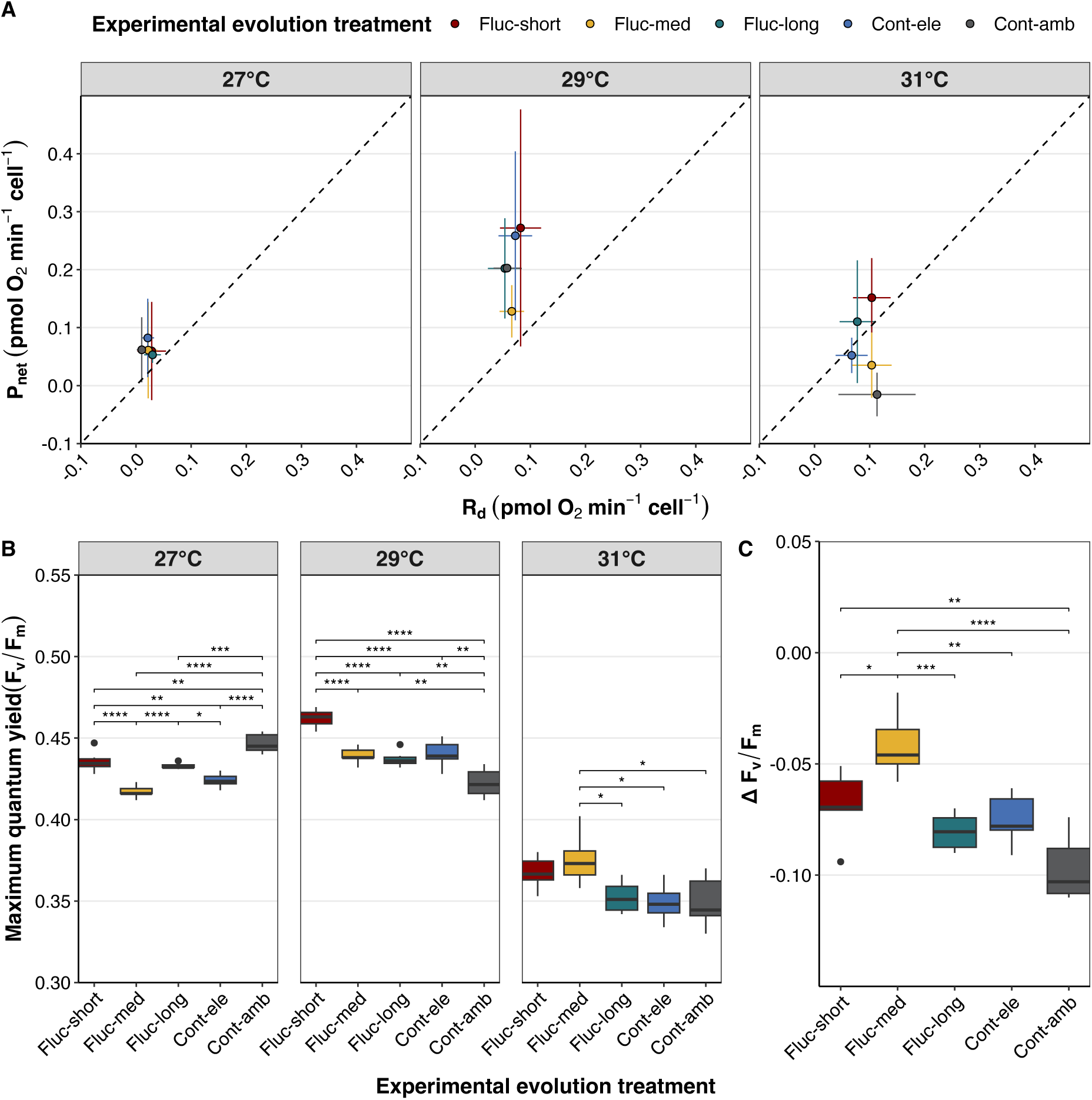
Evolution of net photosynthesis (P_net_) in relation to dark respiration (R_d_, A) and photochemical efficiency (B-C) of *Durusdinium trenchii* lineages under increasing temperatures. (**E**) Deltas in F_v_/F_m_ performance were calculated as the differences between the trait values at 31°C, corresponding to 7 eDHWs of thermal stress, and 27°C. Experimental evolution treatments: Fluc-short = diurnal fluctuations, Fluc-med = fluctuations every 2-3 generations, Fluc-long = fluctuations every 4-5 generations, Cont-ele = continuous exposure to elevated temperatures, Cont-amb = continuous exposure to ambient temperature. Levels of significance: * = p_adj_ < 0.05; ** = p_adj_ < 0.01; *** & **** = p_adj_ < 0.001.

The positive P_gross_ of Fluc-short and to a lesser extent, Fluc-med lineages at 31°C suggests they were able to sustain their energetic demands after 7.2 weeks of experimental degree heating weeks, contrary to the remaining experimental evolution treatments. The increased productivity recorded in the Fluc-short lineages under thermal stress was reflected by an increased F_v_/F_m_ following exposure to elevated temperatures (Fig. 4D; Fig. S13). Although Fluc-med lineages displayed the greatest increase in F_v_/F_m_, relative to Cont-amb, this improvement was not reflected by an increased productivity at 31°C.

#### Melting temperature of the thylakoid membranes does not predict photosystem efficiency, but may explain oxidative stress

Regardless of evolutionary history, pre-exposure to high temperatures (29/31°C) led to an increase in melting temperature (T_m_) of the photosynthetic membranes of all lineages over counterparts grown under ambient (27°C) temperature (Fig. 5A). The temperature-driven difference in T_m_ reflects the plasticity of *D. trenchii*’s photosystem. Increased thermal plasticity was generated in heat-evolved lineages selected under fluctuating conditions, which displayed significant increases (+0.4-0.6°C) relative to Cont-amb (Fig. 5B). Considerable variation in thermal plasticity appeared to have been generated in Fluc-short. The increases in T_m_ were mirrored by lower levels of ExROS accumulating in the culture media as temperatures increased (Fig. 5C,D). Despite the improved thermal plasticity, Fluc-short lineages displayed no changes in levels of ExROS relative to Cont-amb.

**Figure 5:**
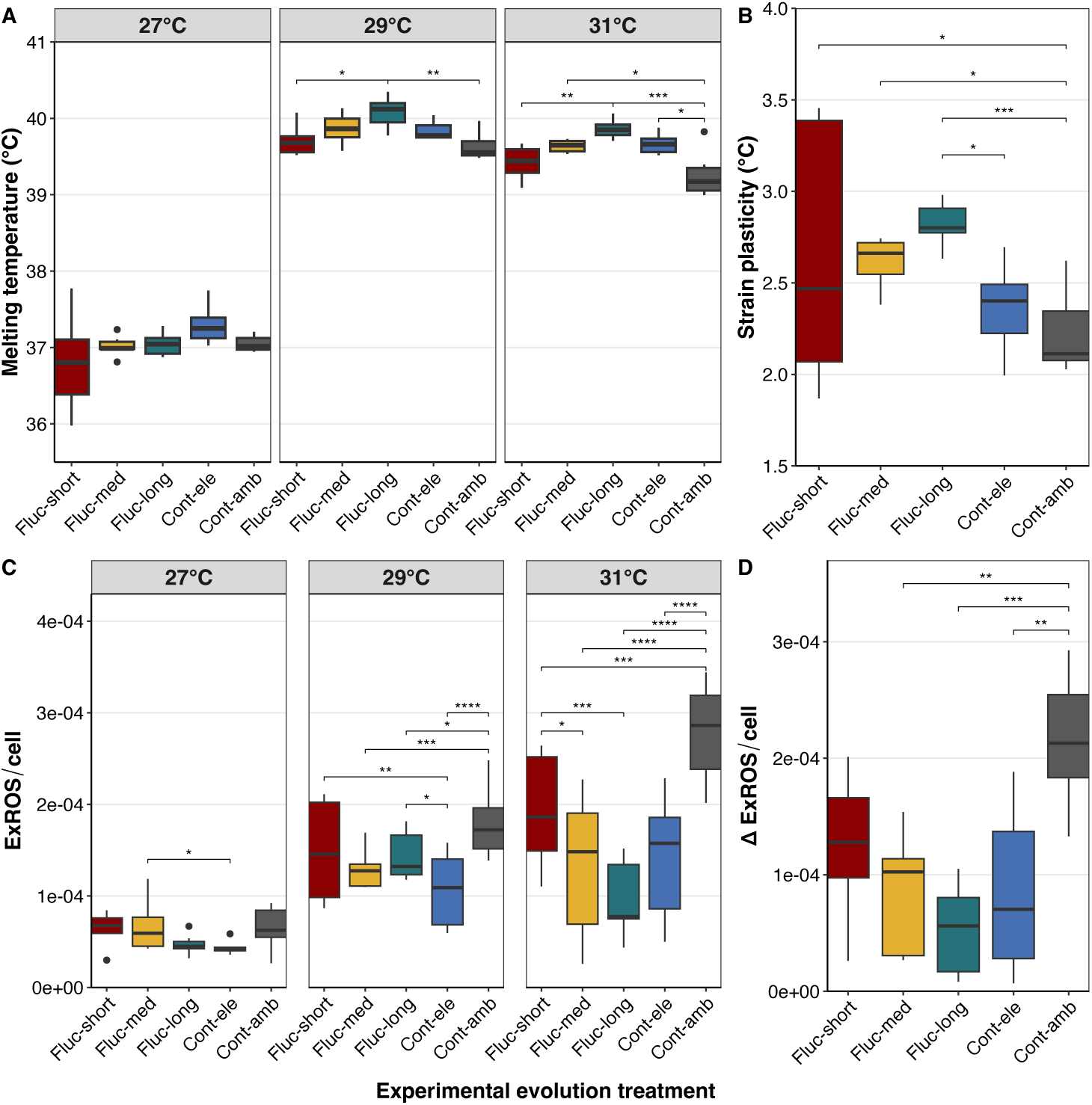
Thylakoid membrane melting temperatures (A) and thermal plasticity (B) of *Durusdinium trenchii* lineages grown under increasing temperatures. Levels of extracellular reactive oxygen species (C-D) measured from culture medium. Lineage-specific melting curves used to obtain the thylakoid membrane melting temperature are available in Fig. S10. (**B**) Thermal plasticity corresponds to the difference in melting temperatures of lineages grown 31 vs. 27 °C. (**D**) Deltas in ExROS levels were calculated as the differences between the trait values at 31°C, corresponding to 7 experimental degree heating weeks of heat stress, and 27°C. Experimental evolution treatments: Fluc-short = diurnal fluctuations, Fluc-med = fluctuations every 2-3 generations, Fluc-long = fluctuations every 4-5 generations, Cont-ele = continuous exposure to elevated temperatures, Cont-amb = continuous exposure to ambient temperature. Levels of significance: * = p_adj_ < 0.05; ** = p_adj_ < 0.01; *** & **** = p_adj_ < 0.001.

#### Trait spaces show greater divergence under thermal stress

We carried out a PCA to understand how the trait responses measured differ between thermal selection strategies (Fig. 6). The PCA illustrates the emergence of different responses to both elevated (31°C) and ambient temperature (27°C), relative to the wild-type lineages, as a result of experimental evolution. Amongst the heat-evolved lineages, responses are unique for Fluc-short, Fluc-med and Cont-ele lineages. The traits photosynthesis (P_net_, T_m_), growth (µ) and oxidative stress (ExROS) drove most of the variation between experimental evolution treatments at 31°C.

**Figure 6:**
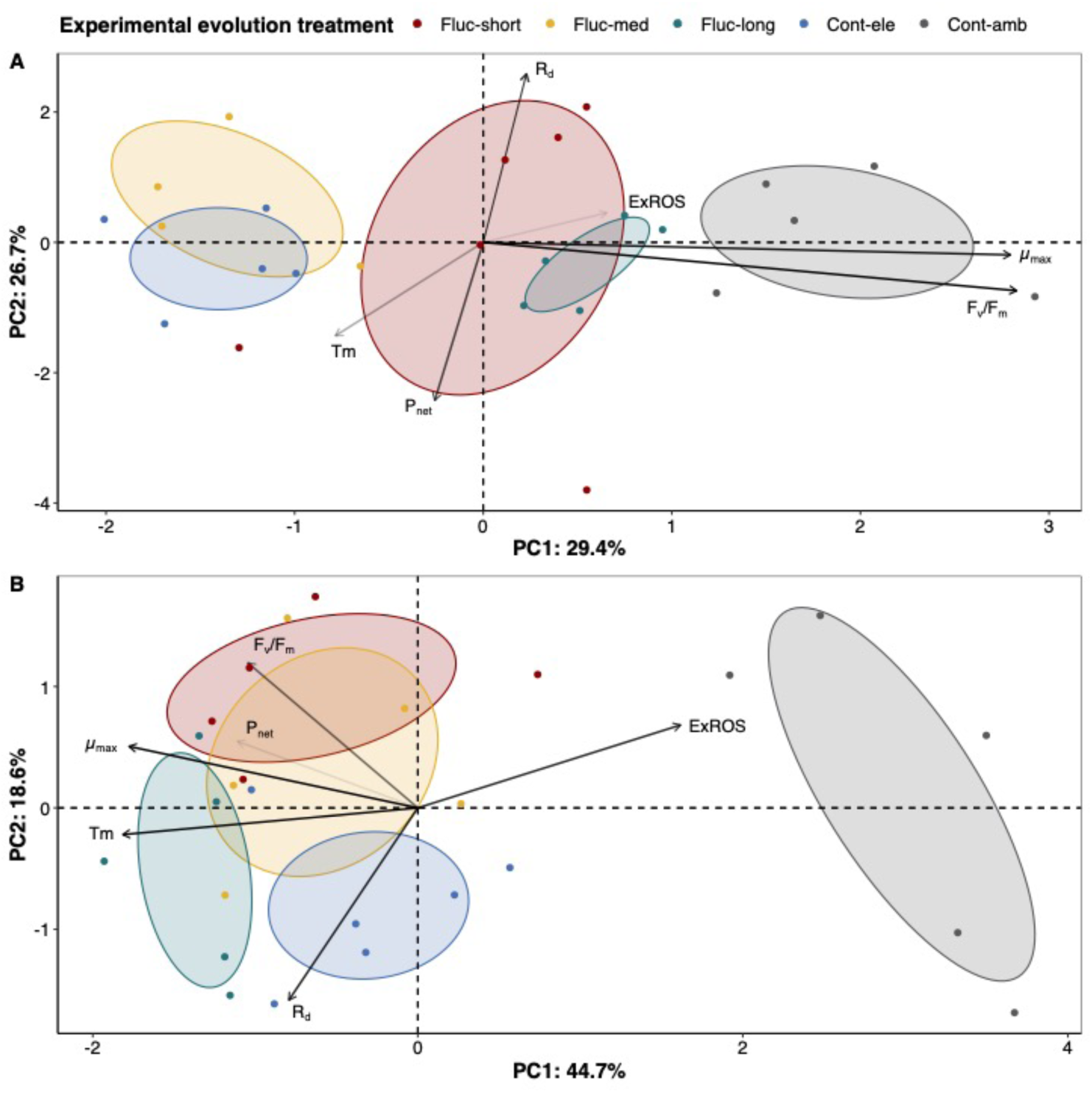
Principal component analyses of trait data collected during the thermal performance assay under ambient (27°C; A) and elevated (31°C; B) temperatures. The weight of each trait’s contribution to the PCA is represented by the shading of the arrows (darker = more contribution, light = less contribution). Ellipses represent 95% confidence intervals. Trait abbreviations: µ = growth rate, F_v_/F_m_ = maximum quantum yield, R_d_ = dark respiration rate, P_net_ = net photosynthesis rate, T_m_ = thylakoid membrane stability. Experimental evolution treatments: Fluc-short = diurnal fluctuations, Fluc-med = fluctuations every 2-3 generations, Fluc-long = fluctuations every 4-5 generations, Cont-ele = continuous exposure to elevated temperatures, Cont-amb = continuous exposure to ambient temperature.

## Discussion

After 2.1 years of experimental evolution, selection under diurnal temperature fluctuations led to the greatest improvement in the thermal niche of *D. trenchii*, driven by increases in both CT_max_ and CT_min_. Our results corroborate reports of daily temperature fluctuations promoting thermal generalism, reflected by an increased T_br_, and that this response correlates with tolerance to novel environments (corresponding to exposure to 21°C in this study) (Bonnefond *et al*., 2017; Ketola *et al*., 2013). However, thermal generalism came with decreased maximum growth rates (r_max_) under optimal temperatures, supporting the view that breadth of adaptation may come at the cost of mean performance in generalists (reviewed by Kassen, 2002). The improved upper thermal tolerance displayed by lineages selected under diurnal fluctuations was associated with an increased photosynthetic capacity (P_net_, P_gross_, F_v_/F_m_, thermal plasticity) under thermal stress, rather than reduced oxidative stress (ExROS).

Evolutionary rescue (i.e., the recovery of a population from extinction as a result of adaptation to environmental stress; Gonzalez *et al*., 2013) likely occurred following initial exposure to 31°C in Fluc-short lineages, as *D. trenchii* lineages were maintained subsequently at temperatures above those that were lethal to the wild-type lineages (≥31°C). Evolutionary studies of the green microalga *Chlamydomonas reinhardtii* reported sexual reproduction to be a beneficial strategy to induce adaptation in short time frames by supporting evolutionary rescue (Bell, 2013; Lachapelle & Bell, 2012; Petkovic & Colegrave, 2019). Given the strong response of Fluc-short lineages to thermal selection after 60-65 generations, sexual reproduction – for which strong evidence now exists in Symbiodiniaceae (Figueroa *et al*., 2021) – may have been key in enabling evolutionary rescue in this study.

Matching the overnight relaxation of temperatures with the dark period may have been an another factor in accelerating adaptation to lethal temperatures. Thermal stress can arrest the progression of mitosis in some symbiodiniaceaens, including *D. trenchii*, in the G_1_ phase, inhibiting cell division which occurs at night (DNA synthesis peaks at the light-to-dark transition time of day) (Fujise *et al*., 2018). Relaxing temperatures overnight could alleviate the negative impact of thermal stress on mitosis, enabling higher rates of asexual propagation. Thus, the strain’s adaptive potential could be additionally improved through the increased occurrence of random somatic mutations.

Selection under fluctuations spanning multiple generations led to minimal fitness (CT_max_) improvements at elevated temperatures. This suggests the relaxation of selective pressure over multiple generations during experimental evolution may limit adaptation to heat in *D. trenchii*. A potential explanation is that the temporary amelioration of conditions in our study reduced the probability of adaptive mutations to spread through the population (Hao *et al*., 2015). Our experimental outcome is in contrast with Schaum *et al*. (2018), who reported adaptation in a marine diatom subjected to fluctuations every 3-4 generations. Beyond species-specific mechanisms, this discrepancy could be due to the difference in the amplitude of the temperature fluctuation between both studies (6°C vs 1-4°C in our study).

Temperature fluctuations over long temporal scales, relative to the *D. trenchii* generation time, did not result in the emergence of thermal generalism. Our results strongly suggest that the frequency of the temperature fluctuation influences adaptive trajectories. Despite limited changes in fitness at elevated temperature, lineages subjected to inter-generational fluctuations displayed improved photosystem plasticity and lower oxidative stress under thermal stress.

Contrary to lineages selected under fluctuating temperatures, continuous selection under elevated temperatures led to changes in thermal performance primarily driven by an increase in T_opt_. The extensive duration (98 weeks) spent at sub-lethal temperatures during experimental evolution likely promoted a greater affinity for warmer temperatures. No evolutionary rescue was observed in Cont-ele lineages once exposed to 31°C, neither before or after recovery at ambient temperature. Nevertheless, the recorded increase in CT_max_ suggests that stable selection under (mostly) sub-lethal temperatures was successful at improving the upper thermal limit of *D. trenchii*. However, the overall improved fitness under warmed conditions came at the cost of a reduced TSM. The decreasing growth rates observed during selection under increasing temperatures suggests this selection strategy may limit the potential for asexual propagation to generate adaptation in the long-term.

We expected that continuous selection under elevated temperatures, relative to the fluctuating treatments, would have led to the emergence of heat-specialists that displayed decreased fitness in lower temperatures. However, our results show that selection at high temperatures also led to a broadening of *D. trenchii*’s thermal niche, suggesting the evolution of a degree of thermal generalism. One hypothesis is that selection under heat may have favoured multiple stress resistance mechanisms, for instance as observed in *Drosophila* (Bubliy & Loeschcke, 2005), which benefited *D. trenchii* at colder temperatures. Continuous thermal selection did not yield any substantial negative adaptive response to thermal stress (only a decrease in TSM), a marked change from previous experimental evolution attempts that used the same *D. trenchii* strain (Chakravarti & van Oppen, 2018; Scharfenstein *et al*., 2023). Beyond the stochastic nature of natural adaptation, a key difference in this study was the long-term exposure to temperatures below 31°C, which may have influenced adaptation. Though no improvements in photosynthetic capacity (P_net_, F_v_/F_m,_ thermal plasticity) were recorded under thermal stress for Cont-ele lineages, reduced oxidative stress was measured. Out of the thermal selection strategies tested, selection under diurnal temperature fluctuations and continuous thermal selection led to the greatest increases in CT_max_ (+0.33°C, σ=0.11 and +0.28°C, σ=0.05, respectively; Fig. 7). From an evolutionary standpoint, the increased thermal tolerance emerged sooner (54-57 generations) in Cont-ele lineages than in Fluc-short lineages (62-65 generations), though rates of adaptation are comparable (∼0.005°C/generation). It appears selection under diurnal temperature fluctuations and continuous thermal selection, primarily at sub-lethal temperatures, are equally viable experimental evolution approaches at increasing Symbiodiniaceae upper thermal tolerance. However, both selection regimes induce unique shifts in thermal performance, promoting either a considerable increase in thermal breadth (Fluc-short) or in thermal optimum (Cont-ele). Our results also suggest that using strategies that vary in the homogeneity of their selective pressure may be valuable to generate lineages presenting differing adaptive mechanisms to heat (e.g., reduced oxidative stress versus improved photosynthetic rates). Exploring whether Fluc-short and Cont-ele lineages can cross-breed (sexually) would be interesting to produce transgressive, extreme phenotypes (Rieseberg *et al*., 1999). Using a community of diverse phenotypes as part of algal symbiont manipulation efforts may increase our ability at generating coral holobionts with improved performance under a range of temperature environments.

**Figure 7:**
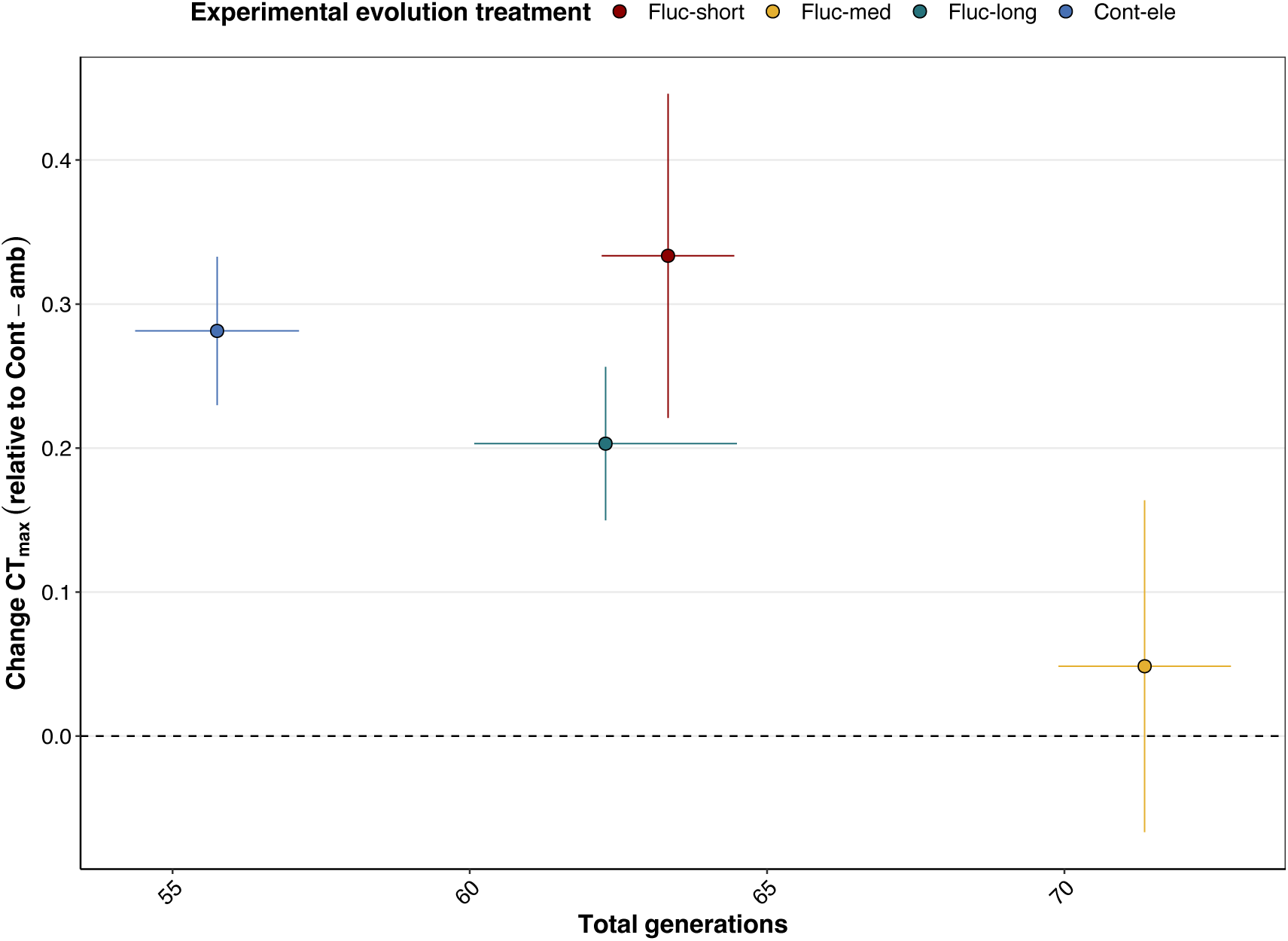
Change in critical thermal maximum (CT_max_) of heat-evolved *D. trenchii* lineages relative to the wild-type lineages as a function of the number of generations elapsed during experimental evolution. Lineranges represent the standard deviation across the experimental evolution treatment. No significant differences in change in CT_max_ were recorded between Fluc-short, Fluc-long and Cont-ele. Experimental evolution treatments: Fluc-short = diurnal fluctuations, Fluc-med = fluctuations every 2-3 generations, Fluc-long = fluctuations every 4-5 generations, Cont-ele = continuous exposure to elevated temperatures, Cont-amb = continuous exposure to ambient temperature.

Additionally, understanding whether certain *in vitro* traits can be indicators of thermal tolerance *in hospite* would be highly valuable. A recent study identified low ROS and high reduced glutathione activity as candidate predictor traits of thermal tolerance *in hospite* (Buerger *et al*., 2023). Given the unique trait spaces generated under thermal stress in some of our heat-evolved lineages, contrasting the performances of these differing phenotypes *in hospite* would be a valuable opportunity to ascertain the role of such traits in coral holobiont thermal tolerance. Further parameterisation of physiological traits in Symbiodiniaceae will provide vital knowledge on the standing functional diversity among algal symbionts and improve our understanding of the mechanisms altered as a result of experimental evolution (Nitschke *et al*., 2022).

In conclusion, our results provide compelling evidence that selection under diurnal temperature oscillations, rather than fluctuations spanning multiple generations, promotes thermal generalism and can improve the fitness of *D. trenchii* at thermal extrema. Continuous selection under elevated temperatures improved fitness under warmed conditions, whilst inducing a degree of thermal generalism. We recommend that future experimental evolution efforts aiming to increase Symbiodiniaceae thermal consider overnight relaxation of temperatures alongside continuous thermal selection, particularly to increase the diversity of phenotypes available for algal symbiont manipulation.

## Acknowledgments

We acknowledge the Bindal people as the Traditional Owners of the land where this research was carried out and pay our respects to Elders past and present. This research was supported by the Australian Research Council Laureate Fellowship to MJHvO (FL180100036), the Paul G. Allen Family Foundation and the Reef Restoration and Adaptation Program, which is funded by the partnership between the Australian Governments Reef Trust and the Great Barrier Reef Foundation. HS and CA thank Toby Wright and Niall Jeeves for their help with the design and building of the chambers used for the respirometry. HS also thanks Dave Hughes for his advice on respiration, Florita Flores for her flow cytometry expertise and Corinne Allen for her help during the experimental evolution.

## Data availability statement

The dataset associated with this study is available in the Supplementary Information and has been archived at the Australian Institute of Marine Science’s Research Data Platform. The dataset can be accessed at: https://apps.aims.gov.au/metadata/view/eceb7313-5c82-40c1-a341-1bf0564c873c. All the code used for this analysis is available at: https://github.com/hscharfenstein/Periodic-and-continuous-thermal-selection-of-D.trenchii.git.

