## Supplementary Information for "Pushing the limits: expanding the temperature tolerance of a coral photosymbiont through differing selection regimes"

### Supplementary material

#### Figures

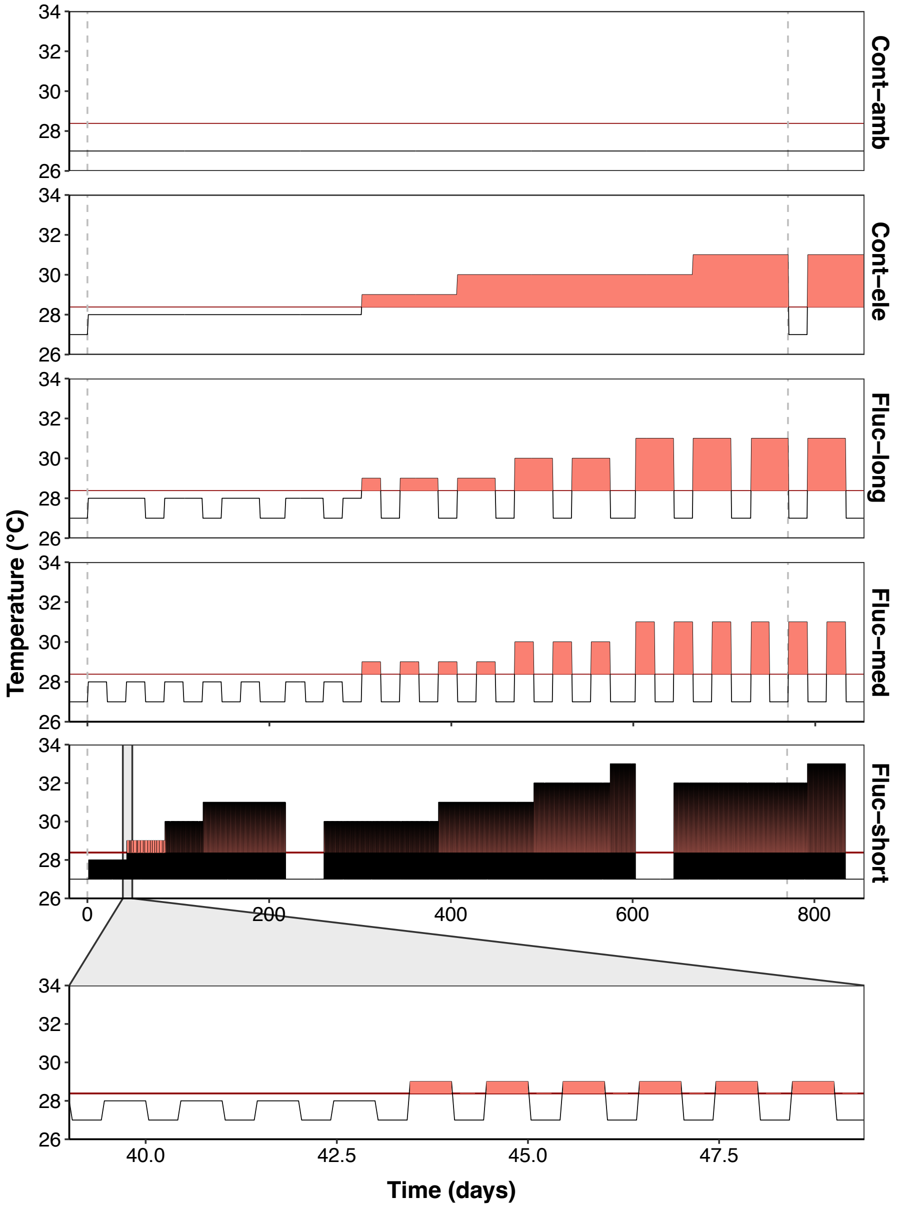


**Figure S1: Temperature profiles used during experimental evolution.** In the case of Fluc-short, ramping up/down between ambient/elevated temperatures was performed over the course of 1 hour. For Fluc-med, Fluc-long and Cont-ele, cultures were transferred between chambers set at the target temperature. The first dashed grey line represents the start of the experimental evolution, whilst the second grey line indicates to the start of the thermal performance assay (t = 770 days/110 weeks). The solid red line reflects the maximum monthly mean climatology for Davies Reef (MMM = 28.38°C), where the strain originated from. Time spent above the MMM is filled in red.

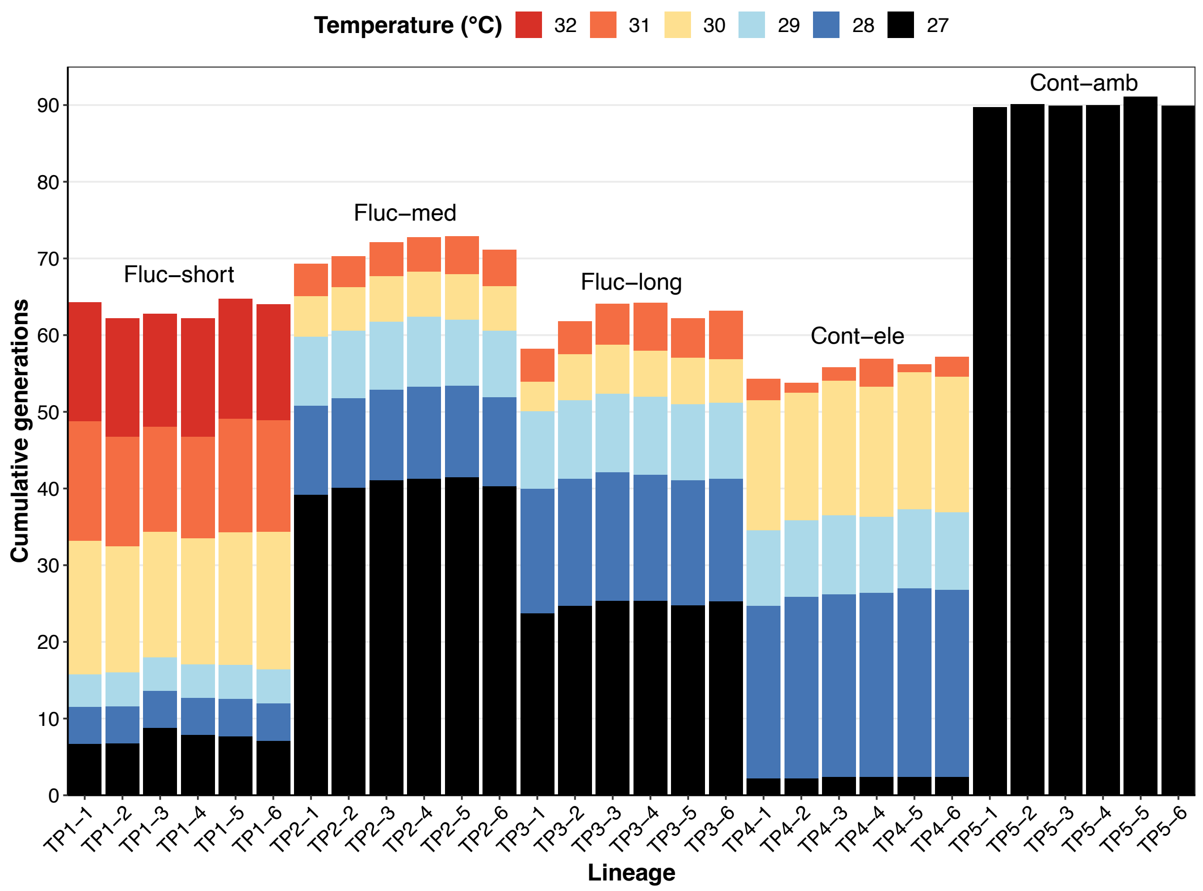


**Figure S2: Number of generations the *Durusdinium trenchii* lineages spent at each temperature during experimental evolution.** In the case of Fluc-short, colours indicate the elevated temperature during the growth cycle. For Fluc-short and Cont-ele, generations spent at 27°C correspond to the unplanned recovery periods following cultures crashing. The number of generations was calculated till the start of the thermal performance assay.

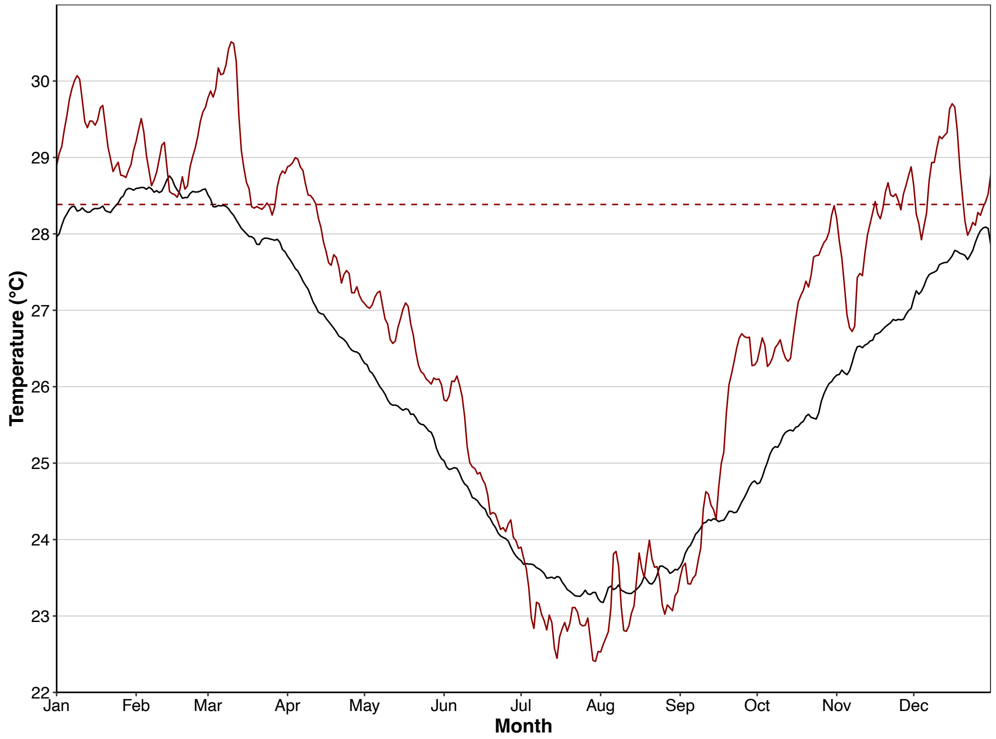


**Figure S3: Daily temperatures recorded at four metres of depth from Davies Reef.** The temperatures were retrieved from the Australian Institute of Marine Science's weather stations (https://weather.aims.gov.au/#/overview). Black line = long-term average (14-16 years) of water temperatures; red line = water temperatures recorded over the course of 2022; dashed red line = maximum monthly mean climatology at Davies Reef (28.38°C).

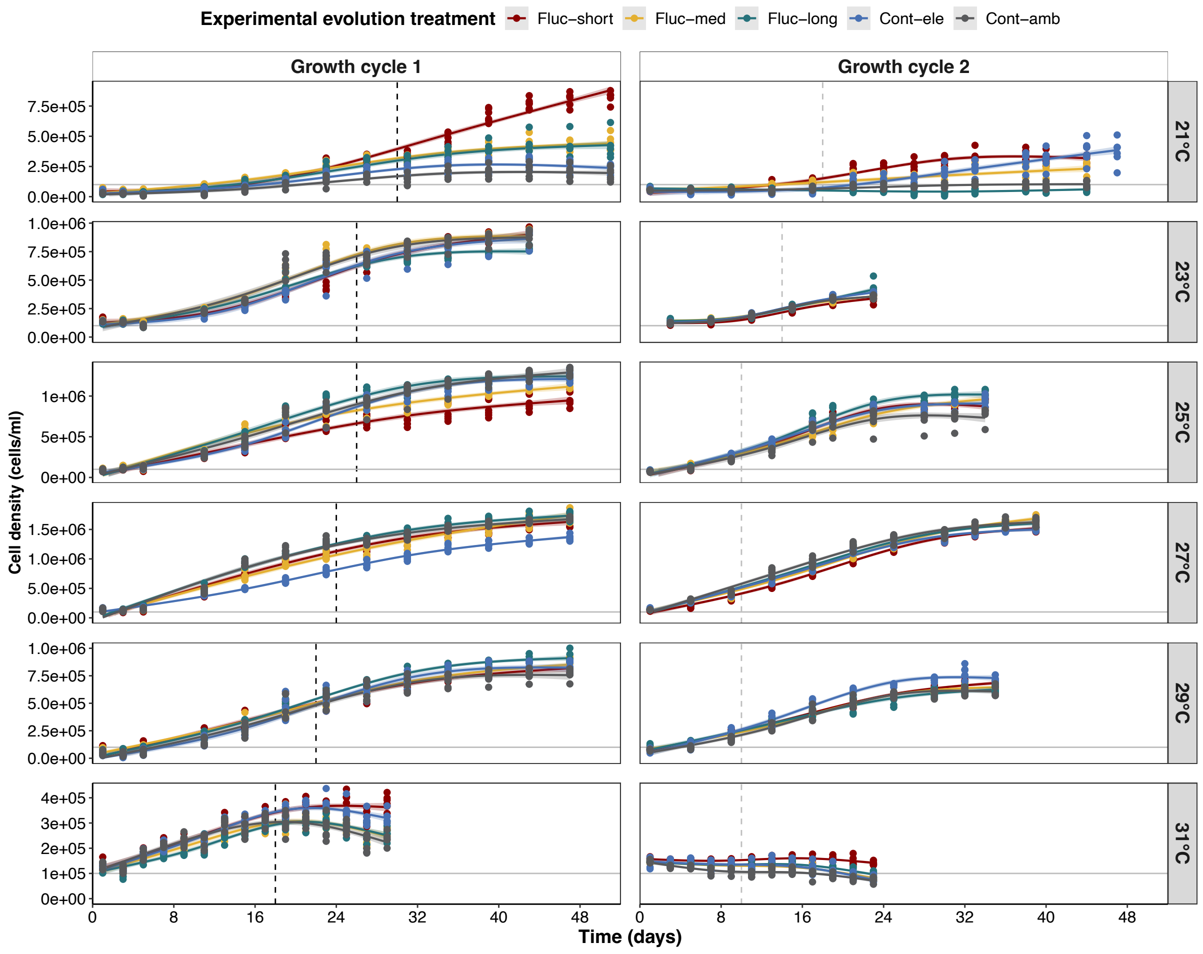


**Figure S4: Growth curves of heat-evolved and wild-type *Duridinium trenchii* lineages during the thermal performance assay.** The black vertical dashed line in growth cycle 1 indicates the timepoint when the lineages were sub-cultured into a fresh culture medium preparation. The grey vertical dashed line in growth cycle 2 indicates the timepoint when trait measurements were performed ($F_v/F_m$, $R_d$, $P_{gross}$, thylakoid membrane $T_m$).

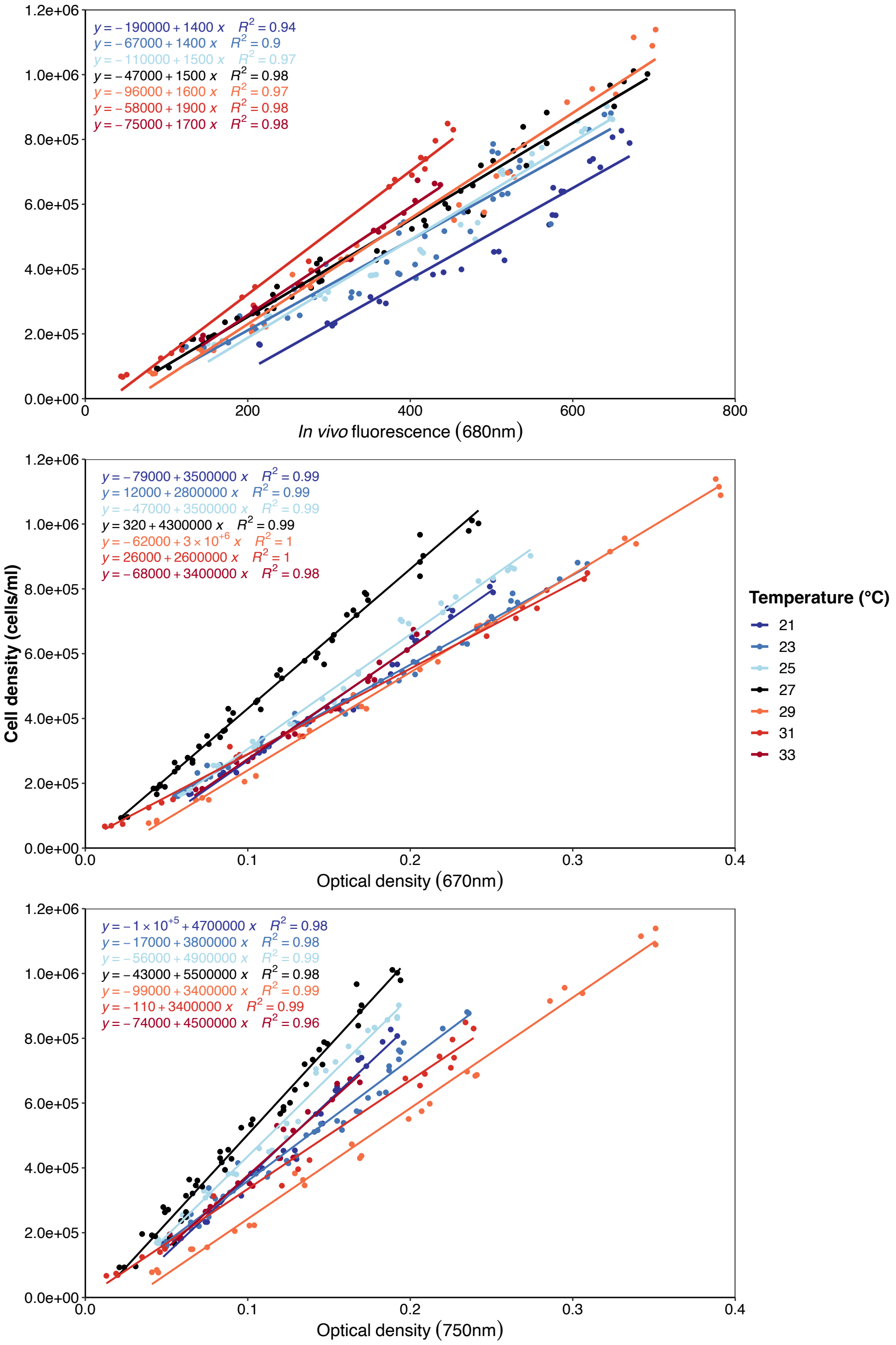


**Figure S5: Calibration curves between *in vivo* fluorescence/optical density measurements and cell densities of wild-type *Durusdinium trenchii* cultures at temperatures ranging 21 to 33°C.** Stock cultures were maintained for 14-16 days (7 days for 33°C) at their growth temperature prior to carrying out measurements to avoid any confounding effects from acclimation on the IVF and OD measurements. Dilutions were prepared from the stock cultures ranging from 1,000,000 to 100,000 cells/ml, with increments of 100,000 cells/ml (n=3 per density). Cell densities were measured via flow cytometry following the same procedure as mentioned in Material and Methods.

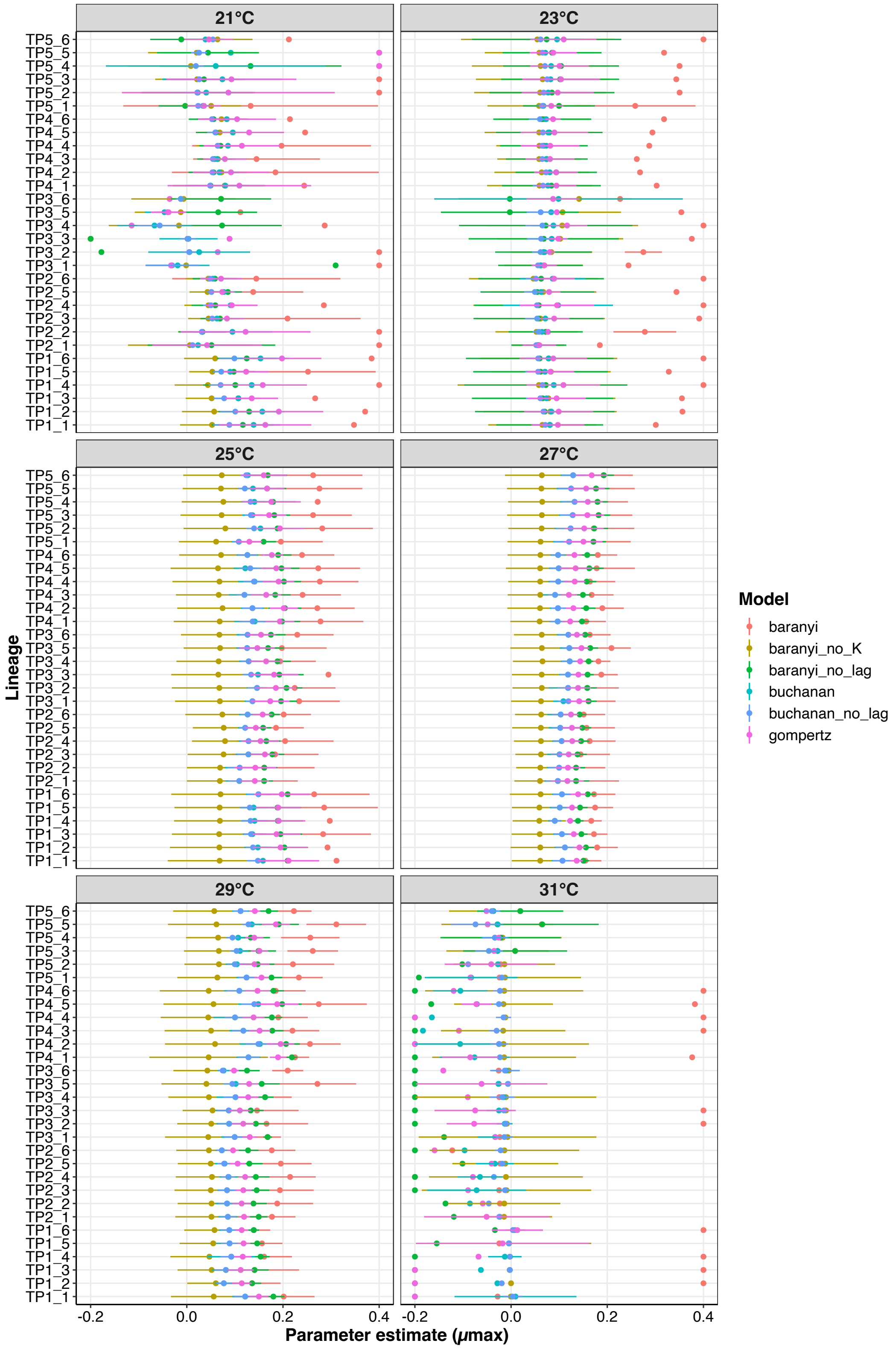


**Figure S6: Maximum growth rate ($\mu_{max}$) estimates obtained from growth model fits.** The parameter estimates obtained from the models *baranyi*, *baranyi_no_lag* and, to a lesser extent, *gompertz* were often found to be at the boundaries of model estimates, particularly at 31°C. Lineranges represent 95% confidence intervals for the parameter estimate.

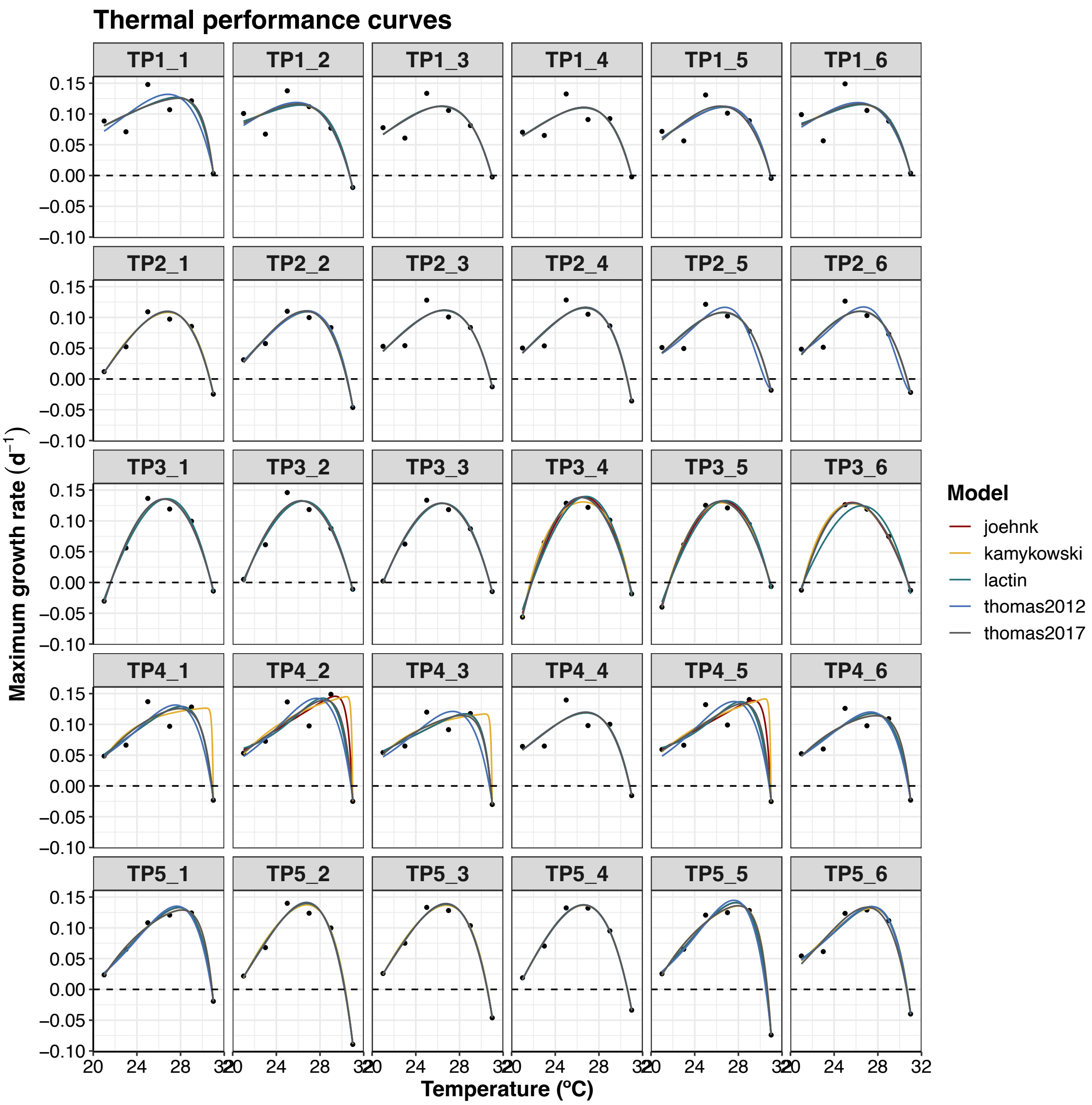


**Figure S7: Lineage specific thermal performance curves generated from the five models tested in this study.**

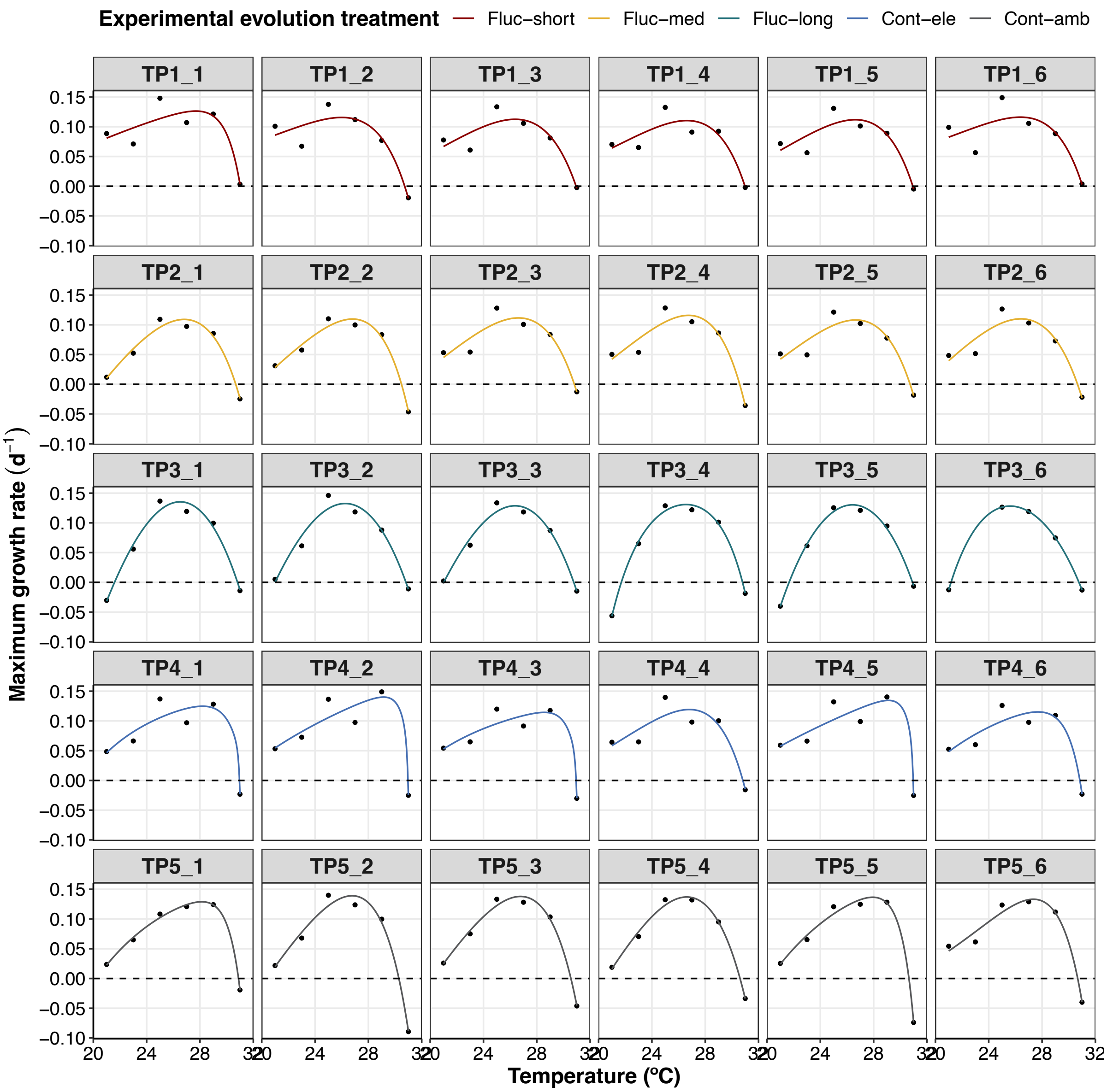


**Figure S8: Average of best performing models.**

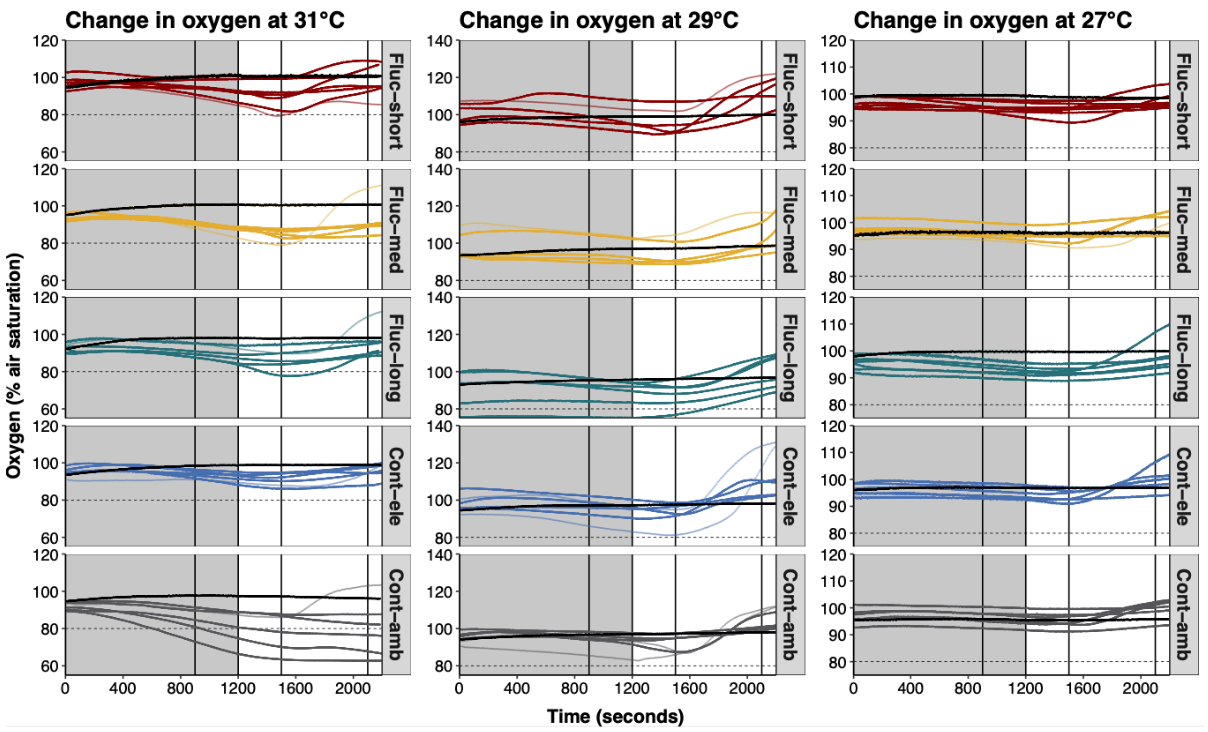


**Figure S9: Oxygen evolution during the dark (shaded) and light phases.** The two vertical lines in the shaded areas indicate the five minute window from which $R_d$ measurements were calculated from. The subsequent two vertical lines indicate the ten minute window in which the $P_{gross}$ measurements were carried out (over a moving sliding window of five minutes selected by *RespR* for best fit). Lines that are faded out indicate samples that were not included in $P_{gross}$ measurements. The horizontal dashed line indicates 80% air saturation, a threshold above which we attempted to maintain oxygen to avoid the possibility of photorespiration. The black lines denote the blank measurements.

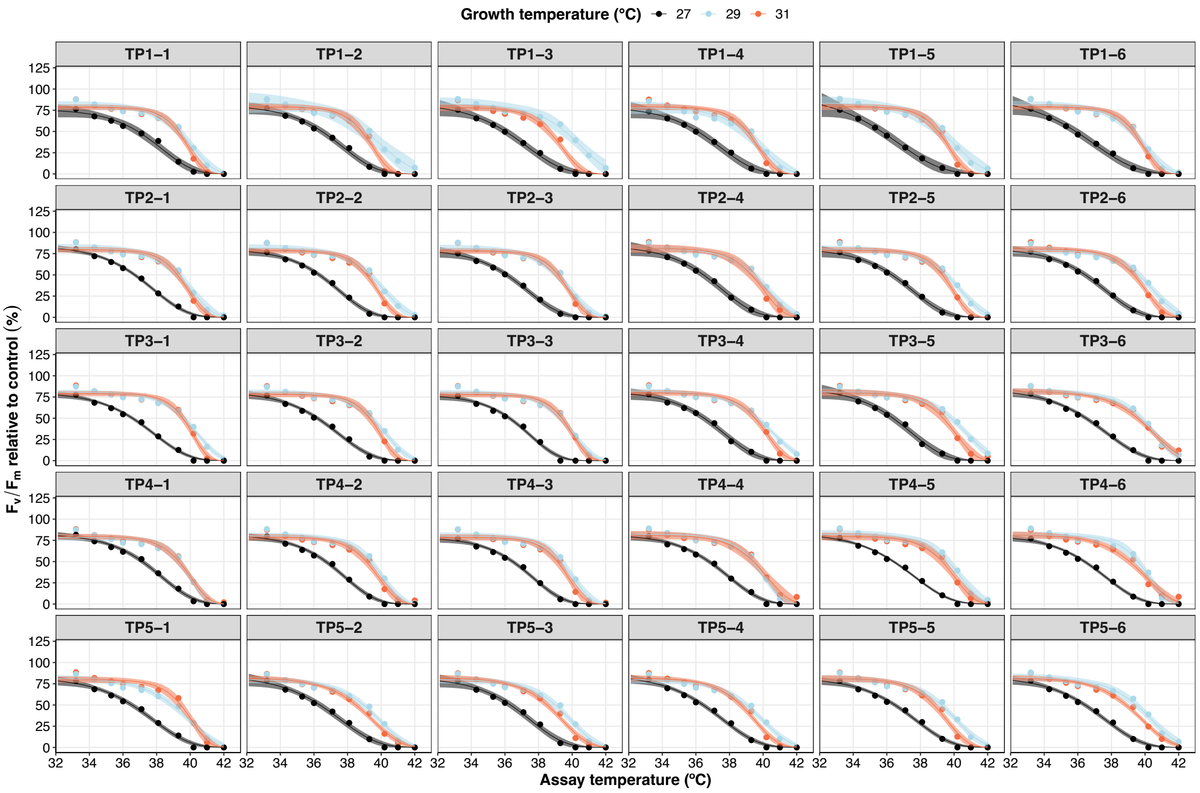


**Figure S10: Temperature-dependent melting curves of thylakoid membranes of *Durusdinium trenchii* lineages grown at 27, 29 and 31°C.** The melting curves represent Weibull model fits. Ribbons reflect the 95% confidence intervals of the model fit across three technical replicates.

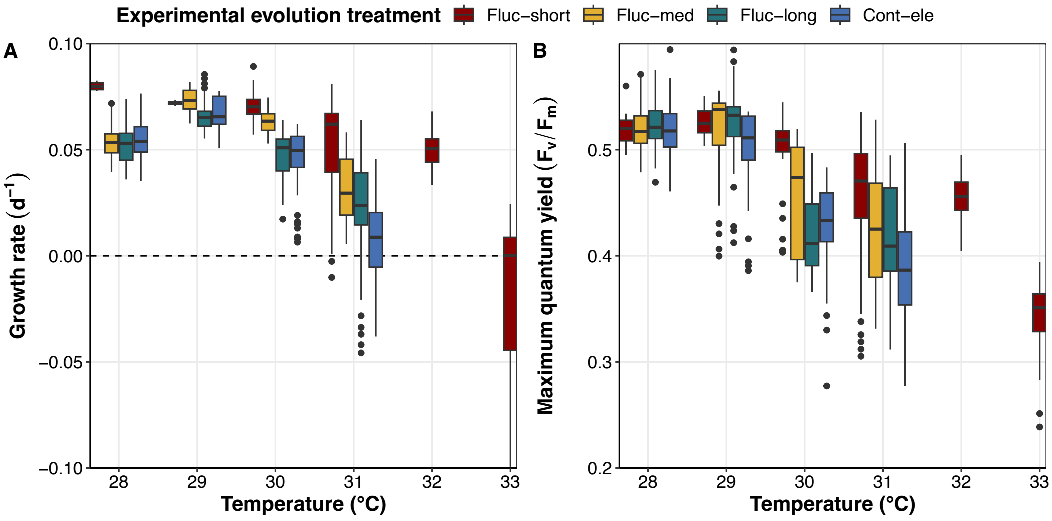


**Figure S11: Growth rates (A) and photochemical efficiencies (B) of heat-evolved *Durusdinium trenchii* lineages at elevated temperatures during experimental evolution.** In the case of Fluc-short, trait performance includes overnight relaxation at ambient temperature.

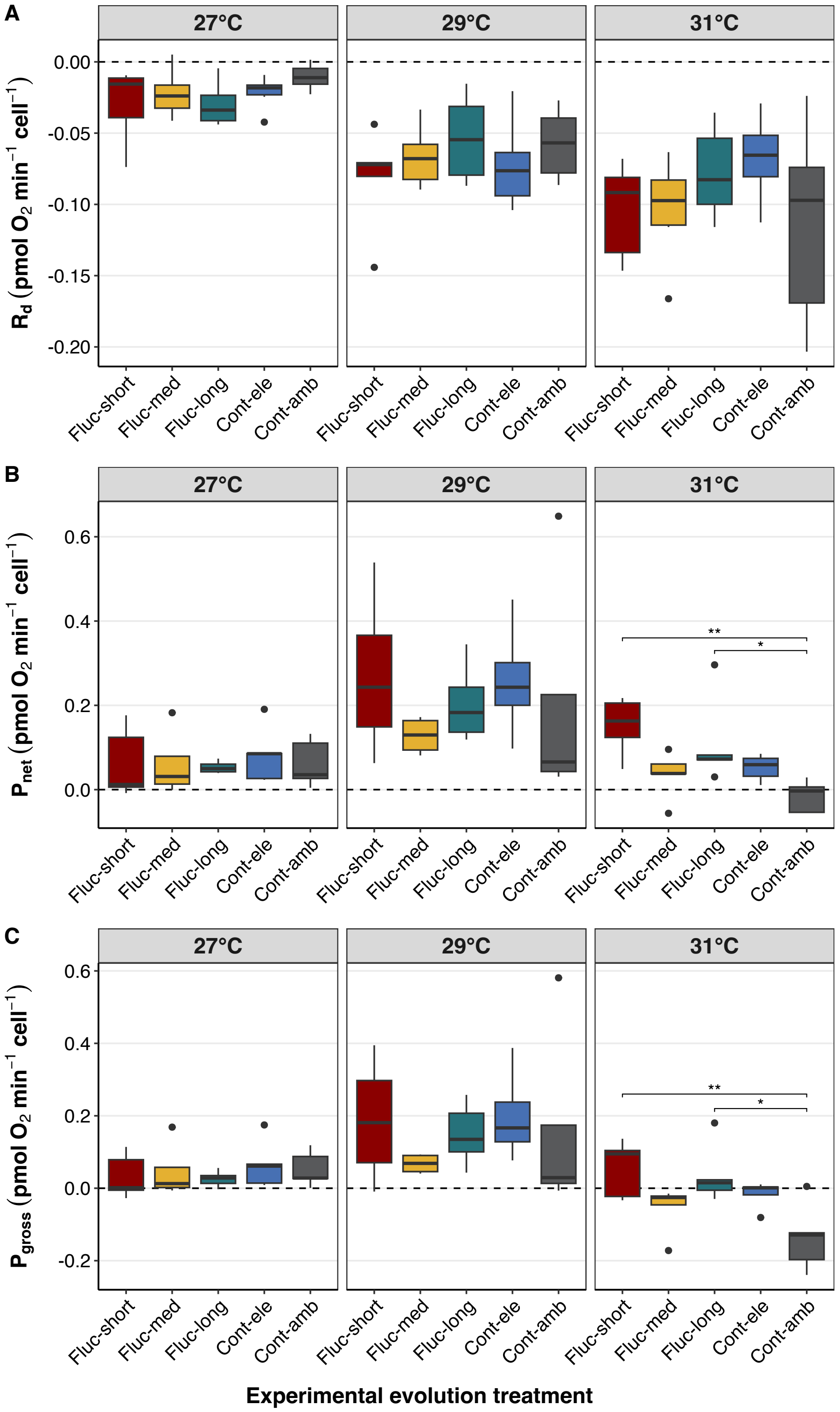


**Figure S12: Respiration (A), net photosynthesis (B) and gross photosynthesis (C) rates of *Durusdinium trenchii* lineages under increasing temperatures.**

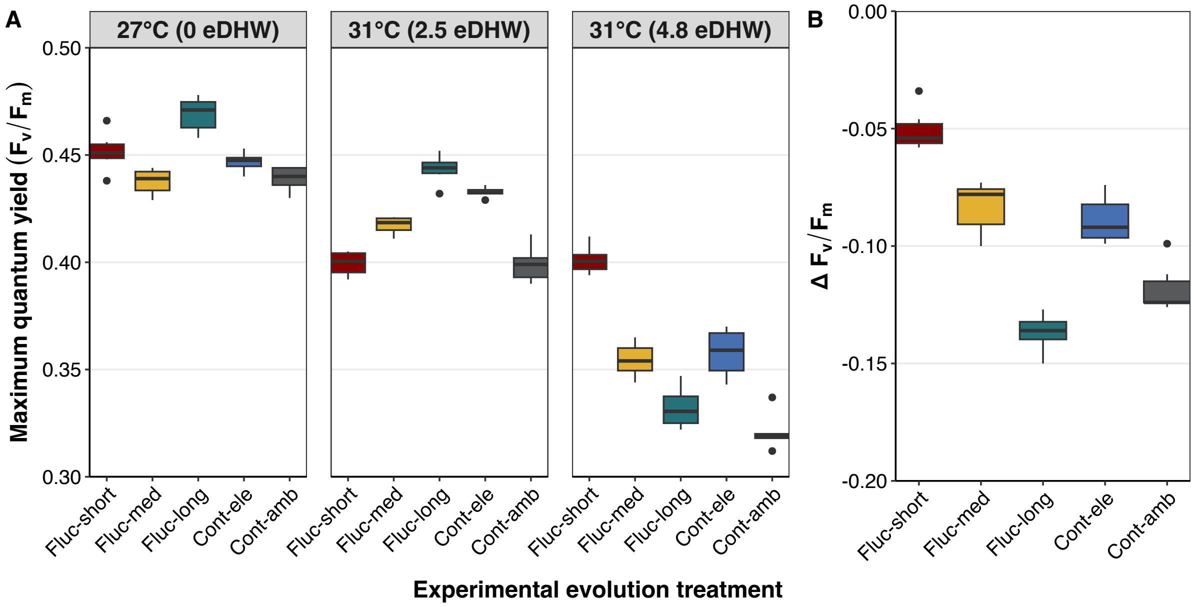


**Figure S13: Photochemical efficiencies of *Durusdinium trenchii* lineages under ambient (27°C) and elevated (31°C) temperatures over the course of the first growth cycle.** (B) Deltas in $F_v/F_m$ levels were calculated as the differences between the trait values at the end of the growth cycle at 31°C, corresponding to 4.8 eDHWs of thermal stress, and the start of the growth cycle at 27°C.

### Tables

| | IMK | Concentrations IMK (mg/l) |
|---|---|---|
| **Water base (MilliQ)** | 1 L | 1 L |
| **Macronutrients (µM)** | | |
| NaNO3 | 2353 | 200 |
| NH4Cl | 50 | 2.68 |
| NaH2PO4 | 9.9 | 1.4 |
| K2HPO4 | 28.7 | 5 |
| Na2EDTA | 110.6 | 37.2 |
| Fe-EDTA | 14.2 | 5.2 |
| Mn-EDTA | 0.96 | 0.332 |
| **Trace Metals (µM)** | | |
| CuSO4 x 5 H2O | 0.01 | 0.0025 |
| ZnSO4 x 7 H2O | 0.08 | 0.023 |
| MnCl2 x 4 H2O | 0.91 | 0.18 |
| Na2MoO4 x 2H2O | 0.03 | 0.0073 |
| CoSO4 x 7 H2O | 0.05 | 0.014 |
| H2SeO3 | 0.01 | 0.0017 |
| **Vitamins (µM)** | | |
| Biotin | 0.0061 | 0.0015 |
| Cyanocobalamin B12 | 0.0011 | 0.0015 |
| Thiamine HCl | 0.593 | 0.2 |

**Table S1: Recipe for the IMK culture medium used in this study.**
